# Normative Modeling of Molecular-Enriched Functional Connectivity for Detecting Deviations from Healthy Brain Aging

**DOI:** 10.64898/2026.09.28.754989

**Authors:** Marco Pinamonti, Manuela Moretto, Leonard Pieperhoff, Prithvi Arunachalam, Frederik Barkhof, Alle Meije Wink, Luigi Lorenzini, Mattia Veronese

## Abstract

**Introduction:** Inter-individual variability in adult brain aging can obscure early pathological alterations and is only partly represented by population-average or scalar brain-aging measures. We combined Receptor-Enriched Analysis of functional Connectivity by Targets (REACT) with normative modeling (NM) to derive spatially resolved, molecularly informed deviation scores for dopamine transporter (DAT)-, norepinephrine transporter (NET)-, and serotonin transporter (SERT)-enriched resting-state functional connectivity (FC). We first evaluated whether these normative models could be transferred to independently processed external data through local calibration and then explored whether the resulting deviation profiles differed with cerebral amyloid burden in cognitively unimpaired older adults.

**Methods:** Hierarchical Bayesian regression (HBR) models with a sinh–arcsinh likelihood (SHASHb) were estimated separately for 204 cortical molecular-enriched FC features in 4,152 healthy adults (18.0–89.8 years) from seven publicly available neuroimaging datasets and evaluated in a held-out healthy test set. Pretrained models were locally adapted to three Amyloid Imaging to Prevent Alzheimer’s Disease Prognostic and Natural History Study (AMYPAD-PNHS) acquisition batches using cognitively unimpaired, amyloid-negative participants (global Clinical Dementia Rating (CDR) = 0; Centiloid (CL) < 10). The independent primary comparison contrasted participants with intermediate amyloid burden (10 ≤ CL < 30; *n* = 111) and amyloid-positive participants (CL ≥ 30; *n*= 66); secondary analyses tested linear associations with continuous CL within participants with CL ≥ 10.

**Results:** Of 204 normative reference models, 199 (97.5%) met the predefined diagnostic criteria; median held-out explained variance (EXPV) was 0.174. Of 612 feature-by-batch transfers, 541 (88.4%) met the strict transfer-diagnostic criteria. In the primary false discovery rate (FDR)-controlled regional analysis, DAT-enriched right caudal middle frontal cortex showed lower locally standardized deviation scores in the intermediate-amyloid-burden group than in the amyloid-positive group (adjusted difference = −0.582, *q* = 0.016). DAT extreme-deviation burden was also greater in the intermediate-amyloid-burden group (difference = 0.032, *q* = 0.044). The right frontal effect was reproduced with model-native scores across the three acquisition batches (pooled standardized effect = −0.698, *q* = 0.010). No NET- or SERT-enriched regional difference between these two groups survived FDR-correction, and no regional or subject-level linear association with continuous CL values survived FDR-correction in the CL ≥ 10 group.

**Conclusion:** NM can provide spatially resolved reference distributions of molecular-enriched FC that are deployable in independently processed external data when local adaptation, calibration, and feature-level transfer diagnostics are incorporated. The AMYPAD-PNHS application identified modest, spatially selective, and molecular-system-specific categorical differences during a cognitively unimpaired stage of amyloid accumulation, while continuous analyses within CL ≥ 10 did not support linear association. These findings support a methodological basis for distributed application of molecular-enriched normative models and motivate independent and longitudinal evaluation of their biological relevance.

**Key Points:**

- Hierarchical normative models characterized 204 regional DAT-, NET-, and SERT-enriched FC features across 4,152 healthy adults; 199/204 models met the predefined normative-reference-model diagnostic criteria.
- Local hierarchical adaptation enabled the pretrained normative models to be applied to independently processed AMYPAD-PNHS data without centralized reprocessing of the target raw images or explicit feature-level harmonization; 541/612 feature–batch transfers met the strict transfer criteria.
- In cognitively unimpaired AMYPAD-PNHS participants, the exploratory amyloid application identified a spatially selective DAT-enriched right frontal difference and greater DAT extreme-deviation burden in the intermediate-amyloid-burden group than in the amyloid-positive group. No FDR-corrected regional NET or SERT differences between these groups and no FDR-corrected linear associations with continuous CL were identified.

## 1 Introduction

Population aging is increasing the prevalence and societal impact of age-related neurodegenerative disorders, including dementia, strengthening the need for imaging approaches capable of characterizing brain alterations before overt clinical impairment (Livingston et al., 2024). A central challenge is the marked heterogeneity of brain aging: individuals of the same chronological age can differ substantially in neural maintenance, reserve, compensation, and vulnerability to pathology (Cabeza et al., 2018; Jylhävä et al., 2017). Conventional population-level analyses summarize this variability at the group level, whereas brain-age approaches reduce high-dimensional neuroimaging information to a single predicted age or brain-age gap (Baecker et al., 2021; Franke & Gaser, 2019). Although useful for global characterization, a single global brain-age estimate provides limited information about where an individual’s brain differs from an age-appropriate reference pattern and which biological systems are preferentially involved (Maccioni et al., 2026; Verdi et al., 2021).

Normative modeling (NM) provides a complementary framework by estimating the conditional distribution of a neuroimaging phenotype within a reference population and expressing an individual’s measurement relative to that distribution (Maccioni et al., 2026; Marquand et al., 2016; Rutherford et al., 2022). This formulation preserves the spatial location, direction, and extremeness of individual deviations rather than requiring abnormalities to be homogeneous across participants. Such individualized reference charts are particularly relevant for neurodegenerative processes, in which individuals with similar clinical or biomarker classifications may show different regional patterns of vulnerability.

Molecular-enriched resting-state functional connectivity (FC) offers a functional phenotype with an additional level of biological context. The blood-oxygenation-level-dependent (BOLD) signal is an indirect hemodynamic correlate of neural activity and does not intrinsically distinguish among neurotransmitter receptors or transporters (Logothetis, 2003, 2008). Receptor-Enriched Analysis of functional Connectivity by Targets (REACT) combines resting-state functional magnetic resonance imaging (fMRI) BOLD fluctuations with population positron-emission tomography (PET)- or single-photon emission computed tomography (SPECT)-derived molecular templates to derive subject-specific FC maps spatially enriched for selected molecular targets (Dipasquale et al., 2019). Using this framework, we previously showed that dopamine transporter (DAT)-, norepinephrine transporter (NET)-, and serotonin transporter (SERT)-enriched FC contains substantial age-related information and that combining the three monoaminergic systems improves age prediction relative to individual systems (Pinamonti et al., 2026). Region-wise NM extends that work from a global prediction problem to an individualized reference-chart framework.

A practical challenge is that large normative reference samples typically combine data acquired across different datasets, scanners, and protocols, while external target data may have been processed through an independently established workflow. Hierarchical Bayesian regression (HBR) is well suited to this setting because acquisition effects can be represented hierarchically rather than removed from the data, and pretrained models can subsequently be adapted to previously unseen batches using local reference observations (Bayer et al., 2022; Kia et al., 2022; Rutherford et al., 2022). The use of a flexible sinh–arcsinh likelihood (SHASHb) additionally allows conditional location, scale, skewness, and tail behavior to be represented for non-Gaussian neuroimaging phenotypes (A. A. A. de Boer et al., 2024). This creates a deployment-oriented scenario in which a normative model can be distributed to a new hospital or imaging center and calibrated locally, provided that the downstream phenotype definition remains compatible. In the present study, the target Amyloid Imaging to Prevent Alzheimer’s Disease Prognostic and Natural History Study (AMYPAD-PNHS) data had been preprocessed independently of the seven-dataset normative reference cohort; we therefore preserved a common downstream definition while using local hierarchical adaptation, calibration, and transfer diagnostics to address the unseen acquisition environment.

The AMYPAD-PNHS provides a biologically relevant setting in which to evaluate this framework. Contemporary criteria conceptualize Alzheimer’s disease (AD) as a biological process that can begin before measurable cognitive decline, with amyloid PET serving as a core in-vivo biomarker of amyloid pathology (Jack et al., 2024). Functional alterations have been reported during cognitively unimpaired and preclinical stages, including changes in default-mode activity and connectivity (Ingala et al., 2021; Sperling et al., 2009), and these changes need not vary monotonically with amyloid burden: phases of hyperconnectivity and hypoconnectivity have been described across early amyloid and tau accumulation, and inverted U-shaped associations between amyloid and default-mode connectivity have been reported in cognitively normal older adults (Palmqvist et al., 2017; Schultz et al., 2017; Wu et al., 2026). Within AMYPAD-PNHS, amyloid burden has also been associated with altered structure–function coupling in older adults without dementia (Arunachalam et al., 2026). Although region-wise NM of molecular-enriched REACT phenotypes has been demonstrated in psychiatric populations (Lawn et al., 2024), and REACT has been applied to AD cohorts with partial amyloid characterization (Manca et al., 2025), to our knowledge molecular-enriched normative models have not been transferred to an independent amyloid PET-characterized cohort to quantify individual deviations. A cognitively unimpaired, amyloid-characterized cohort therefore provides an informative setting in which to test whether REACT-based normative models can identify molecularly and spatially specific functional alterations preceding measurable cognitive decline.

Accordingly, this study had two objectives. First, we tested whether regional age- and sex-conditioned distributions of DAT-, NET-, and SERT-enriched FC could be estimated in a large healthy adult reference cohort, validated in held-out healthy participants, and locally transferred to three previously unseen AMYPAD-PNHS acquisition batches. We hypothesized that local HBR adaptation would provide adequately calibrated deviation scores for most cortical molecular-enriched features, while feature-level diagnostics would identify features remaining sensitive to target-domain shift. Second, we examined the transferred deviation profiles of participants at risk of developing dementia, who were currently experiencing no measurable cognitive decline. These participants were referred to as cognitively unimpaired and were defined as having a global Clinical Dementia Rating (CDR) score of 0. They spanned the amyloid-negative, intermediate-amyloid-burden, and amyloid-positive groups. We tested subject-level burden, regional signed deviations and extreme-deviation prevalence, the independent difference between the intermediate-amyloid-burden and amyloid-positive groups, linear associations with continuous Centiloid (CL) within CL ≥ 10, and potential moderation by apolipoprotein E (APOE) *ɛ*4 carrier status. This exploratory application was intended to determine whether the transferred normative representation could reveal spatially and molecularly specific functional differences associated with amyloid burden during a clinically presymptomatic stage.

## 2 Methods

### 2.1 Study design and ethics

The present study employed a two-stage NM design, comprising model estimation and validation in a large, multi-site healthy normative reference cohort, followed by model transfer and application to an independent target cohort from the AMYPAD-PNHS.

The healthy normative reference cohort comprised de-identified data from publicly accessible neuroimaging datasets. Each contributing study obtained approval from its respective local research ethics committee or institutional review board, and all participants provided informed consent according to the procedures of the original studies.

The AMYPAD-PNHS (EudraCT 2018-002277-22) is a multicenter European study of older adults without dementia (Pieperhoff et al., 2026). The study was approved by the Medical Ethics Review Committee of Amsterdam UMC under application number 2018.357, along with the relevant local ethical approvals from the contributing parent cohorts. All participants provided written informed consent. The present analyses employed de-identified data procured under the existing AMYPAD-PNHS ethical approvals and data-use agreements.

### 2.2 Study cohorts

#### 2.2.1 Healthy normative reference cohort

The healthy normative reference cohort was assembled from seven publicly available neuroimaging datasets spanning the adult lifespan: the Cambridge Centre for Ageing and Neuroscience (Cam-CAN) (Shafto et al., 2014; Taylor et al., 2017), the Dallas Lifespan Brain Study (DLBS) (Park et al., 2025), the Lifespan Human Connectome Project in Aging (HCP-Aging) (Bookheimer et al., 2019; Harms et al., 2018), the Human Connectome Project Young Adult (HCP-YA) (Smith et al., 2013; Van Essen et al., 2013), the UCLA Consortium for Neuropsychiatric Phenomics (LA5c) (Poldrack et al., 2016), the Nathan Kline Institute Rockland Sample (NKI-RS) (Nooner et al., 2012), and the Southwest University Adult Lifespan Dataset (SALD) (Wei et al., 2018).

For the present study, only participants classified as healthy according to the recruitment and screening procedures of the original source studies were retained in the normative cohort. Across datasets, the source protocols excluded major neurological and psychiatric disorders and applied study-specific health, cognitive, substance-use, and magnetic resonance imaging (MRI)-safety eligibility criteria appropriate to the corresponding cohort (Bookheimer et al., 2019; Nooner et al., 2012; Park et al., 2025; Poldrack et al., 2016; Shafto et al., 2014; Van Essen et al., 2013; Wei et al., 2018). For LA5c, only participants belonging to the healthy-control subgroup were included (Poldrack et al., 2016). Participants were further retained when the demographic variables and imaging-derived features required for NM were available and the corresponding T1-weighted (T1w) and resting-state functional MRI (rs-fMRI) data satisfied the study-specific imaging quality control (QC) criteria described in Supplementary Section A.1.1. For HCP-YA, only the first available rs-fMRI run was retained for each participant. Participants older than 90 years were excluded to restrict model estimation to the predefined adult age range. The resulting analysis-ready sample was therefore treated throughout the study as a healthy normative reference cohort for estimating expected age- and sex-conditioned variation in molecular-enriched FC. Its demographic characteristics are reported in Section 3.1.

### MRI acquisition

The MRI acquisition protocols exhibited variation across the normative datasets with respect to scanner platform, sequence parameters, spatial resolution, and resting-state acquisition duration. A summary of dataset-specific acquisition parameters for T1w and rs-fMRI is provided in Tables 1 and 2, respectively.

**Table 1:** T1-weighted structural MRI acquisition parameters across the normative reference datasets.

| Dataset | Scanner | Sequence | TR<br>[ms] | TE<br>[ms] | TI<br>[ms] | FA<br>[°] | FOV<br>[mm] | Voxel size<br>[mm] |
| --- | --- | --- | --- | --- | --- | --- | --- | --- |
| Cam-CAN | 3T Siemens TIM Trio | MPRAGE | 2250 | 2.99 | 900 | 9 | 256 × 240 × 192 | 1 × 1 × 1 |
| DLBS | 3T Philips Achieva | MPRAGE | 8.1 | 3.7 | 1100 | 12 | 204 × 256 × 160 | 1 × 1 × 1 |
| HCP-Aging | 3T Siemens Prisma | ME-MPRAGE | 2500 | 1.8/3.6/5.4/7.2 | 1000 | 8 | 256 × 240 × 166 | 0.8 × 0.8 × 0.8 |
| HCP-YA | 3T Siemens Connectome Skyra | MPRAGE | 2400 | 2.14 | 1000 | 8 | 224 × 224 | 0.7 × 0.7 × 0.7 |
| LA5c | 3T Siemens Trio | MPRAGE | 2530 | 3.31 | 1100 | 7 | 256 × 256 × 176 | 1 × 1 × 1 |
| NKI-RS | 3T Siemens TIM Trio | MPRAGE | 1900 | 2.52 | 900 | 9 | 256 × 246 × 176 | 1 × 1 × 1 |
| SALD | 3T Siemens Trio | MPRAGE | 1900 | 2.52 | 900 | 9 | 256 × 256 × 176 | 1 × 1 × 1 |

*Note.* MPRAGE = magnetization-prepared rapid gradient echo; ME-MPRAGE = multi-echo magnetization-prepared rapid gradient echo; TR = repetition time; TE = echo time; TI = inversion time; FA = flip angle; FOV = field of view. Sources: Cam-CAN (Shafto et al., 2014; Taylor et al., 2017); DLBS (Park et al., 2025); HCP-Aging (Bookheimer et al., 2019; Harms et al., 2018); HCP-YA (Van Essen et al., 2013); LA5c (Poldrack et al., 2016); NKI-RS (Nooner et al., 2012); SALD (Wei et al., 2018).

**Table 2:**
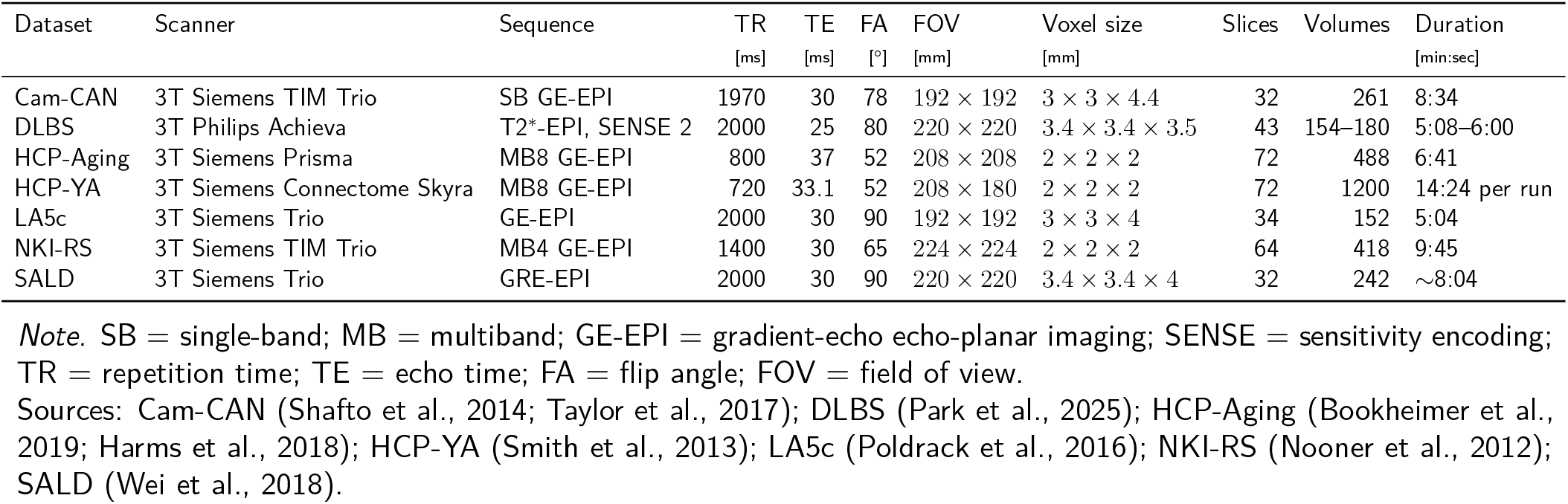
Resting-state functional MRI acquisition parameters across the normative reference datasets.

| Dataset | Scanner | Sequence | TR<br>[ms] | TE<br>[ms] | FA<br>[°] | FOV<br>[mm] | Voxel size<br>[mm] | Slices | Volumes | Duration<br>[min:sec] |
| --- | --- | --- | --- | --- | --- | --- | --- | --- | --- | --- |
| Cam-CAN | 3T Siemens TIM Trio | SB GE-EPI | 1970 | 30 | 78 | 192 × 192 | 3 × 3 × 4.4 | 32 | 261 | 8:34 |
| DLBS | 3T Philips Achieva | T2*-EPI, SENSE 2 | 2000 | 25 | 80 | 220 × 220 | 3.4 × 3.4 × 3.5 | 43 | 154–180 | 5:08–6:00 |
| HCP-Aging | 3T Siemens Prisma | MB8 GE-EPI | 800 | 37 | 52 | 208 × 208 | 2 × 2 × 2 | 72 | 488 | 6:41 |
| HCP-YA | 3T Siemens Connectome Skyra | MB8 GE-EPI | 720 | 33.1 | 52 | 208 × 180 | 2 × 2 × 2 | 72 | 1200 | 14:24 per run |
| LA5c | 3T Siemens Trio | GE-EPI | 2000 | 30 | 90 | 192 × 192 | 3 × 3 × 4 | 34 | 152 | 5:04 |
| NKI-RS | 3T Siemens TIM Trio | MB4 GE-EPI | 1400 | 30 | 65 | 224 × 224 | 2 × 2 × 2 | 64 | 418 | 9:45 |
| SALD | 3T Siemens Trio | GRE-EPI | 2000 | 30 | 90 | 220 × 220 | 3.4 × 3.4 × 4 | 32 | 242 | ~8:04 |

*Note.* SB = single-band; MB = multiband; GE-EPI = gradient-echo echo-planar imaging; SENSE = sensitivity encoding; TR = repetition time; TE = echo time; FA = flip angle; FOV = field of view.
Sources: Cam-CAN (Shafito et al., 2014; Taylor et al., 2017); DLBS (Park et al., 2025); HCP-Aging (Bookheimer et al., 2019; Harms et al., 2018); HCP-YA (Smith et al., 2013); LA5c (Poldrack et al., 2016); NKI-RS (Nooner et al., 2012); SALD (Wei et al., 2018).

**Table 3:**
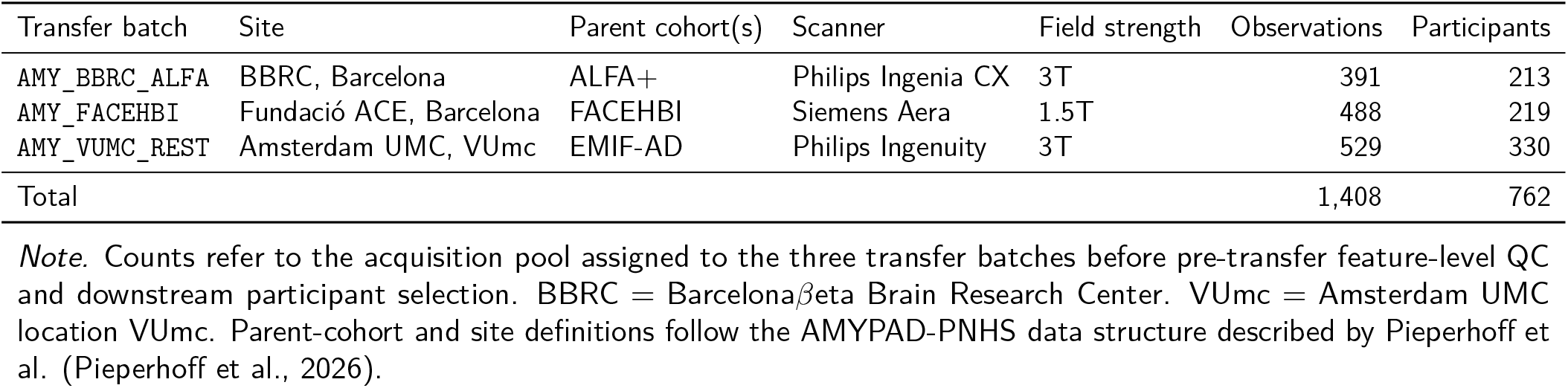
AMYPAD-PNHS acquisition groups included in the normative-model transfer workflow.

| Transfer batch | Site | Parent cohort(s) | Scanner | Field strength | Observations | Participants |
| --- | --- | --- | --- | --- | --- | --- |
| AMY_BBRC_ALFA | BBRC, Barcelona | ALFA+ | Philips Ingenia CX | 3T | 391 | 213 |
| AMY_FACEHBI | Fundació ACE, Barcelona | FACEHBI | Siemens Aera | 1.5T | 488 | 219 |
| AMY_VUMC_REST | Amsterdam UMC, VUmc | EMIF-AD | Philips Ingenuity | 3T | 529 | 330 |
| Total |  |  |  |  | 1,408 | 762 |

*Note.* Counts refer to the acquisition pool assigned to the three transfer batches before pre-transfer feature-level QC and downstream participant selection. BBRC = Barcelona $\beta$ eta Brain Research Center. VUmc = Amsterdam UMC location VUmc. Parent-cohort and site definitions follow the AMYPAD-PNHS data structure described by Pieperhoff et al. (Pieperhoff et al., 2026).

#### 2.2.2 AMYPAD-PNHS target cohort

The target cohort was derived from the AMYPAD-PNHS, a multicenter European cohort designed to investigate amyloid-*β* pathology and related imaging and clinical biomarkers in older adults without dementia (Pieperhoff et al., 2026). The AMYPAD-PNHS MRI resource encompasses structural and rs-fMRI in conjunction with amyloid PET, demographic, clinical, cognitive, and genetic information that has been collected across multiple European parent cohorts and study sites. The principal AMYPAD-PNHS eligibility criteria relevant to the present study were age above 50 years, absence of dementia as indicated by a global CDR score below 1, and the ability to undergo MRI and amyloid PET examinations (Pieperhoff et al., 2026).

For the present study, participants were required to have rs-fMRI and T1w images and demographic variables necessary for the molecular-enriched FC and normative-model transfer workflows. For analyses related to amyloid, an available amyloid PET measurement was also required. MRI and amyloid PET acquisitions were required to be separated by no more than 365 days. When multiple resting-state acquisitions were available for a participant, only the first rs-fMRI session satisfying this temporal criterion was retained; consequently, each participant contributed a single rs-fMRI observation to the transfer and downstream analytical workflow. Acquisition groups were retained for NM transfer when at least 30 unique local reference participants were available. Local reference participants were defined as cognitively unimpaired, amyloid-negative individuals, operationally defined as having global CDR = 0 and cerebral amyloid burden below 10 CL. Three acquisition groups satisfied these requirements: ALFA+ acquired at the Barcelona*β*eta Brain Research Center (BBRC), FACEHBI acquired at Fundació ACE, and data acquired at Amsterdam UMC location VUmc. These groups were represented in the normative-model transfer workflow as AMY_BBRC_ALFA, AMY_FACEHBI, and AMY_VUMC_REST, respectively.

Prior to the implementation of feature-level QC and downstream participant selection, the three acquisition groups comprised 1,408 rs-fMRI observations from 762 unique participants. The ALFA+ group comprised 391 observations from 213 participants, FACEHBI comprised 488 observations from 219 participants, and VUmc comprised 529 observations from 330 participants. The final participant numbers following the implementation of QC measures, the availability of clinical and amyloid measures, and downstream sample selection are reported in Section 3.2.

### MRI acquisition

The acquisition of MRI data in AMYPAD-PNHS exhibited heterogeneity across parent cohorts and sites, thereby reflecting the multicenter nature of the study (Pieperhoff et al., 2026). For the present analysis, the target dataset was restricted to the three transfer groups previously described: ALFA+ at the Barcelona*β*eta Brain Research Center, FACEHBI at Fundació ACE, and data acquired at Amsterdam UMC location VUmc. The principal T1w and rs-fMRI acquisition characteristics of the included protocols are summarized in Tables 4 and 5, respectively. For further details, please refer to the work of Pieperhoff et al. (2026).

**Table 4:** T1-weighted structural MRI acquisition parameters for the AMYPAD-PNHS acquisition protocols represented in the present study.

| Site/cohort | Scanner | TR<br>[ms] | TE<br>[ms] | FA<br>[°] | Slice thickness<br>[mm] |
| --- | --- | --- | --- | --- | --- |
| ALFA+ | 3T Philips Ingenia CX | 9.9 | 4.6 | 8 | 0.75 |
| FACEHBI | 1.5T Siemens Aera | 2200 | 2.7 | 8 | 1.0 |
| VUmc | 3T Philips Ingenuity | 8 | 4.5 | 8 | 1.0 |

*Note.* TR = repetition time; TE = echo time; FA = flip angle.

**Table 5:** Resting-state functional MRI acquisition parameters for the AMYPAD-PNHS acquisition protocols represented in the present study.

| Site/cohort | Scanner | TR<br>[ms] | TE<br>[ms] | FA<br>[°] | Slice thickness<br>[mm] | Time points | Duration<br>[min:sec] |
| --- | --- | --- | --- | --- | --- | --- | --- |
| ALFA+ | 3T Philips Ingenia CX | 1600 | 35 | 70 | 3.1 | 300 | 8:00 |
| FACEHBI | 1.5T Siemens Aera | 4000 | 50 | 90 | 3.5 | 150 | 10:00 |
| VUmc | 3T Philips Ingenuity | 1800 | 35 | 80 | 3.0 | 202 | 6:04 |

*Note.* TR = repetition time; TE = echo time; FA = flip angle.

### Amyloid-PET acquisition and Centiloid quantification

The amyloid PET measurements provided within AMYPAD-PNHS were used to assess cerebral amyloid-*β* burden. The acquisition and standardization procedures have been previously delineated for AMYPAD-PNHS (Arunachalam et al., 2026). Briefly, PET scans were acquired 90–110 min after administration of either [^18^F]flutemetamol at 185 MBq (±10%) or [^18^F]florbetaben at 350 MBq (±20%), using tracer-specific protocols comprising four 5-min frames. The processing of PET images was conducted using the automated AMYPAD pipeline, an automated framework that incorporates frame-wise motion correction, averaging, and co-registration to the corresponding structural MRI. Standardized uptake value ratio images were calculated in native space using the whole cerebellum as the reference region. Subsequently, the global cortical amyloid burden was expressed on the CL scale using the Global Alzheimer’s Association Interactive Network standard target region (Arunachalam et al., 2026).

CL values were employed in two distinct ways. First, they were used to define the local reference participants required for normative-model transfer. Second, they were used to characterize amyloid burden in downstream analyses, as detailed in Sections 2.5.3 and 2.6.

### 2.3 MRI preprocessing and quality control

The healthy normative-reference cohort was processed using the same overall workflow adopted in our previous molecular-enriched brain-age study, comprising the sequential use of MRI Quality Control (MRIQC), functional MRI Preprocessing (fMRIPrep), eXtensible Connectivity Pipeline for DCAN (XCP-D), and FMRIB Software Library (FSL) (Pinamonti et al., 2026). For the present study, the principal preprocessing change was the use of a 0.01 Hz temporal high-pass cutoff in XCP-D, rather than the 0.001 Hz cutoff used previously. The AMYPAD-PNHS rs-fMRI data had been independently preprocessed using the established workflow described by Arunachalam et al. (2026) and were not retrospectively reprocessed through the complete normative-reference pipeline; this deliberate upstream mismatch formed part of the external-transfer evaluation. The main upstream differences from the reference workflow were the spatial smoothing kernel (4 versus 6 mm full-width at half-maximum (FWHM)), temporal filtering (0.01–0.1 Hz band-pass versus 0.01 Hz high-pass), and the nuisance model (six motion parameters with mean white matter (WM) and cerebrospinal fluid (CSF) signals, without censoring, versus motion parameters with their derivatives and anatomical CompCor components with interpolation of high-motion frames). Full quality-control and preprocessing details for both cohorts are reported in Supplementary Section A.1.1.

### 2.4 Molecular-enriched functional connectivity

Molecular-enriched FC was derived with REACT using the same two-stage framework and the same DAT, NET, and SERT molecular templates used in our previous brain-age study (Pinamonti et al., 2026). As in that study, the three templates were supplied together as a single four-dimensional molecular atlas and modeled simultaneously in both REACT stages, so that each molecular-enriched map was estimated conditionally on the other two molecular regressors; accordingly, the Stage-1 analysis mask was restricted to voxels in which all three templates had positive values. The principal feature-definition change was the use of the subject-specific Desikan–Killiany (DK) cortical parcellation (Desikan et al., 2006): mean molecular-enriched FC was extracted from 68 cortical regions per molecular system, yielding 204 cortical molecular-system features, whereas the previous study used a broader Schaefer/Harvard–Oxford/SUIT regional feature set. The same molecular templates, analysis masks, DK definitions, and regional extraction procedure were applied to the normative-reference and AMYPAD-PNHS cohorts to preserve downstream feature correspondence for model transfer. Full REACT, molecular-template, and regional feature-extraction details are reported in Supplementary Section A.1.2.

### 2.5 Normative modeling

#### 2.5.1 Model specification

The estimation of NMs was conducted using hierarchical Bayesian regression (HBR), implemented in Predictive Clinical Neuroscience toolkit (PCNtoolkit) (v1.3.0) (Bayer et al., 2022; S. de Boer et al., 2026; Rutherford et al., 2022). The analysis encompassed one model for each of the 204 cortical molecular-enriched FC features; these models are hereafter referred to as the normative reference models. In the analysis, age and sex were included as biological covariates. The acquisition dataset was represented as a hierarchical batch effect to model systematic differences among the seven reference imaging cohorts. The numerical coding scheme assigned female = 0 and male = 1. Prior to estimation, model covariates and response variables underwent standardization within the PCNtoolkit modeling framework.

The SHASHb was employed to accommodate potential non-Gaussianity in the conditional response distribution (A. A. A. de Boer et al., 2024; Jones & Pewsey, 2009). The likelihood function is defined by the following parameters: conditional location (*µ*), scale (*σ*), skewness (*ɛ*), and tail weight (*δ*). The model for *µ* and *σ* incorporated a hierarchical random intercept for the acquisition dataset, with both *µ* and *σ* being modeled as functions of age and sex. Covariate effects were shared across datasets, whereas dataset-specific intercepts allowed the conditional location and scale to differ among acquisition cohorts. The shape parameters, denoted by the variables *ɛ* and *δ*, were estimated as common parameters across datasets.

The non-linear age effects were represented by an identifiable cubic B-spline basis with five knots. The constant direction previously represented by the model intercept was eliminated from the standard B-spline representation. The remaining basis components were then centered and orthogonalized without rescaling. This process yielded six linearly independent spline basis functions while maintaining the underlying cubic-spline function space. Sex was maintained as a linear covariate in the statistical analysis. The resulting design matrix was subjected to automatic full column rank verification prior to model fitting.

Weakly informative priors were specified on the standardized scale. For *µ*, regression coefficients were assigned N (0, 10) priors, while dataset-specific intercepts followed a centered hierarchical prior with population-level location N (0, 1) and a positive hierarchical scale obtained through a soft-plus transformation of a N (0, 1) prior. For *σ*, regression coefficients followed N (0, 2) priors, while hierarchical dataset intercepts were centered on a N (1, 1) population-level prior with a positive soft-plus-transformed scale derived from a N (0, 1) prior. A soft-plus mapping was additionally applied to *σ* to enforce positivity. The skewness parameter *ɛ* followed a N (0, 1) prior, whereas the positive tail-weight parameter *δ* was obtained through a soft-plus transformation of a N (1, 1) prior.

#### 2.5.2 Model estimation and internal validation

The healthy normative reference cohort was subdivided at the participant level into independent training and held-out test sets using an 80/20 split with a fixed random seed of 42. In order to preserve both the representation of the dataset and the age distributions, dataset-specific age bins were generated using quantile-based binning. The dataset was split into groups based on both the dataset and the age bin. The final seven-dataset healthy reference cohort comprised 3,324 participants in the training set and 828 participants in the held-out healthy test set, with no participant represented in both subsets.

The estimation of model parameters was conducted exclusively from the training data. Posterior sampling was performed with the No-U-Turn Sampler (NUTS) (Hoffman & Gelman, 2014) implemented in PyMC (Abril-Pla et al., 2023) through PCNtoolkit. For each cortical feature, four independent Markov chains were executed, with 1,000 tuning iterations followed by 1,500 posterior draws per chain. This process yielded a total of 6,000 post-warm-up draws. The sampling method employed utilized the jitter+adapt_full initialization, a target acceptance probability of 0.95, a maximum tree depth of 12, and a fixed sampling seed.

After estimation, each fitted model was applied without refitting to the corresponding held-out observations. Predictive performance was summarized from the PCNtoolkit test-set statistics, with explained variance (EXPV), standardized mean squared error (SMSE), and mean standardized log loss (MSLL) treated as the principal predictive and probabilistic performance measures. *R* ^2^ and root-mean-squared error (RMSE) were retained as complementary descriptive metrics when available. The calibration was characterized independently from the held-out normative-deviation scores by their mean, standard deviation (SD), and maximum absolute magnitude.

Posterior and sampler diagnostics were retained for every fitted cortical model, including the potential scale reduction statistic (*R*^^^) (Vehtari et al., 2021), bulk and tail effective sample size (ESS), Monte Carlo standard errors (MCSEs), divergent transitions, maximum-tree-depth saturation, and Bayesian fraction of missing information (BFMI). A predefined diagnostic screen required maximum *R*^^^ ≤ 1.01, minimum bulk ESS ≥ 400, minimum tail ESS ≥ 400, no divergent transitions, a fraction of iterations reaching maximum tree depth ≤ 0.01, and minimum BFMI ≥ 0.30. Held-out deviation scores were additionally required to be finite for all available test observations, to have absolute mean ≤ 0.50, SD between 0.50 and 1.50, and maximum absolute value ≤ 20.

#### 2.5.3 Transfer to the AMYPAD-PNHS cohort

The fitted reference models were transferred separately to three predefined AMYPAD acquisition batches (AMY_BBRC_ALFA, AMY_FACEHBI, and AMY_VUMC_REST) using the PCNtoolkit transfer framework. Transfer and local calibration were performed independently within each acquisition batch.

Prior to transfer, an observation-level QC procedure was applied to the regional REACT matrix within each acquisition batch. For each feature, robust standardized values were calculated using the within-batch median and median absolute deviation, with interquartile-range or standard-deviation fallbacks when required. An observation was automatically excluded when at least 50% of evaluable regional features had an absolute robust standardized value > 10, when the median absolute robust standardized value exceeded 25, when more than 20% of regional values were missing, or when fewer than 100 regional features were evaluable.

Local transfer-reference participants were cognitively unimpaired, amyloid-negative individuals, with CDR = 0 and CL < 10. Unique local reference participants were randomly partitioned using a deterministic feature- and batch-specific seed into approximately 80% adaptation and 20% held-out validation subsets, subject to a minimum of 20 adaptation participants and six held-out validation participants. Batches were eligible for transfer only when at least 30 local reference participants were available.

Transfer was performed in two stages. First, the reference model was adapted using only the adaptation subset and then evaluated in the untouched local held-out subset. The transferred model-native deviation score was denoted *Z*^raw^. To assess local transfer calibration independently of the held-out observations, held-out raw deviation scores were centered and scaled using the mean and SD estimated only from the adaptation subset.

A feature–batch transfer was considered to pass the transfer diagnostics when two conditions were met. First, the adapted model had to satisfy the posterior and sampler criteria used for the normative reference models (Section 2.5.2). Second, the held-out locally standardized deviation scores had to be finite and to show adequate calibration, defined as an absolute mean ≤ 0.10, an SD between 0.80 and 1.20, and a maximum absolute value ≤ 5. Because each held-out subset contained approximately 20% of the local reference participants (minimum six), these criteria screen for gross miscalibration rather than establishing calibration precisely.

After this validation step, a final transfer was performed for each feature and acquisition batch using all available unique local reference participants. Transfer used the same HBR-SHASHb structure and age basis learned during normative-model estimation, with PCNtoolkit transfer freedom set to 1.0. Posterior sampling used four chains, 500 tuning iterations, and 1,500 posterior draws per chain, with PyMC NUTS sampling, jitter+adapt_full initialization, target acceptance probability 0.95, maximum tree depth 12, and fixed sampling seed. The final adapted model was then used to generate predictions and deviation scores for every complete retained AMYPAD participant in the corresponding batch, with one selected rs-fMRI session per participant.

Two deviation-score representations were retained. The model-native score, *Z*^raw^, was the deviation returned directly by the transferred HBR-SHASHb model. For cross-batch primary analyses, a locally calibrated score was additionally calculated for each feature *r* and acquisition batch *b* as

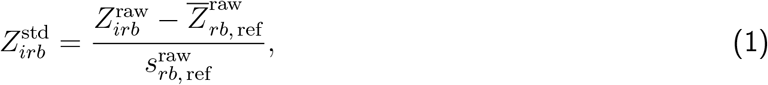

where 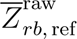 and 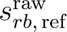 are the mean and SD of *Z*^raw^ among all unique local reference participants used in the final adaptation for that feature–batch combination. This score is denoted *Z*^std^. It was used as the primary deviation representation because it places transferred scores on a common local reference scale across acquisition batches; *Z*^raw^ was retained for batch-stratified robustness analyses.

#### 2.5.4 Individual cortical deviation measures

For each participant, positive deviation scores indicated molecular-enriched FC that exceeded the age-, sex-, and batch-conditioned normative expectation, whereas negative scores indicated values that fell below that expectation (Maccioni et al., 2026; Rutherford et al., 2022). The primary AMYPAD feature domain was fixed a priori at the complete set of 204 cortical molecular-system features.

Regional analyses were conducted on a feature-by-feature basis, with all complete cases utilized for the variables stipulated by the corresponding model. Subject-level multiregional burden measures necessitate a comprehensive finite deviation profile within the pertinent predefined domain (all 204 cortical features for the global measure, or the 68 features belonging to the relevant molecular system for system-specific measures).

### Extreme-deviation burden

For the primary locally calibrated representation, a regional observation was classified as extreme when |*Z*^std^| > 1.96. For subject *i* and a predefined feature domain containing *R* regions, extreme-deviation burden was expressed as the proportion

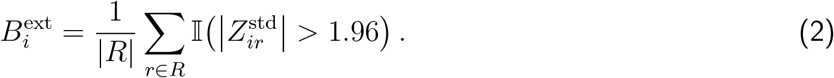

The burden was calculated on a global scale across the 204 cortical features and separately across the 68 DAT, 68 NET, and 68 SERT features. Positive and negative extreme counts were retained as descriptive audit measures; the absolute extreme-deviation proportion was the extreme-burden measure used for inference.

### Ranked absolute deviation burden

To retain information about the full multi-regional deviation profile rather than thresholding deviations dichotomously, ranked absolute deviation burden (RADB) was calculated for each participant. Within a predefined feature domain containing *R* regions, absolute deviation magnitudes were ordered from largest to smallest,

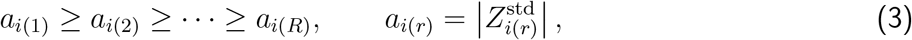

and plotted against *x_r_* = log_10_(*r*). The trapezoidal area under this ranked deviation curve was calculated as

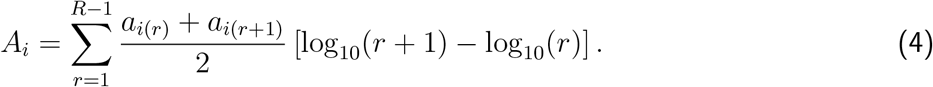

To remove the trivial dependence of the area on the length of the log-rank axis, the area was normalized by the corresponding log-rank range,

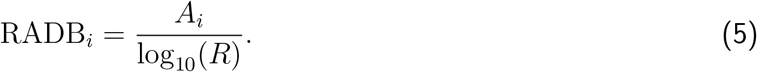

Thus, RADB summarizes the magnitude of a participant’s distributed cortical normative deviations, giving greater influence to the largest deviations while incorporating the entire ordered profile. RADB was calculated globally and separately for DAT, NET, and SERT from *Z*^std^. Normalization by log_10_(*R*) removes the trivial dependence on the length of the log-rank axis; however, because the distribution of ordered deviations also depends on the number and correlation structure of features within a domain, RADB values were interpreted within each predefined feature scope rather than compared directly across scopes containing different numbers of features.

### 2.6 Statistical analysis

All amyloid-related analyses were restricted to cognitively unimpaired participants, operationally defined in the present study as having a global CDR score of 0. A single baseline observation per participant was selected before construction of the analytical dataset, and participant uniqueness was verified.

Participants were categorized as amyloid-negative (CL < 10), intermediate amyloid burden (10 ≤ CL < 30), and amyloid-positive (CL ≥ 30). These operational categories follow the AMYPAD CL context-of-use framework, in which values below 10 exclude amyloid pathology with high certainty, values above 30 correspond well with pathological amyloid amounts, and values between these cutoffs define an intermediate range associated with an increased risk of disease progression (Collij et al., 2024). Amyloid-negative participants contributed to local model transfer and to feature- and batch-specific calibration of *Z*^std^; comparisons involving this group were therefore interpreted as reference-anchored analyses rather than fully independent case–control tests. The predefined independent categorical comparison was between the intermediate-amyloid-burden and amyloid-positive groups.

All tests were two-sided. Ordinary least squares (OLS) models were estimated with heteroscedasticity-consistent covariance estimator (HC3) standard errors (SEs) (MacKinnon & White, 1985). Unless explicitly stated otherwise, adjusted cross-batch regression models included age, sex, and AMYPAD acquisition batch. The Benjamini–Hochberg false discovery rate (FDR) procedure (Benjamini & Hochberg, 1995) was applied within predefined inferential families, with *q* < 0.05 considered statistically significant. For primary regional analyses, multiplicity correction used the fixed family of 204 cortical molecular-system features. For global and molecular-system-specific subject-level analyses, multiplicity correction was performed across the four predefined scopes (global, DAT, NET, and SERT) separately for each outcome and analytical question.

#### 2.6.1 Global and system-specific deviation burden

Overall evidence for differences across the amyloid-negative, intermediate-amyloid-burden, and amyloid-positive groups was assessed using Welch one-way tests, followed by all three pairwise Welch comparisons.

For the predefined independent comparison, RADB was additionally analyzed using HC3 regression contrasting the intermediate and amyloid-positive groups while adjusting for age, sex, and acquisition batch. RADB was the primary subject-level outcome. Extreme-deviation burden was evaluated as a complementary threshold-based subject-level outcome using the same adjusted intermediate-versus-positive framework. For each burden outcome, the global, DAT, NET, and SERT models constituted a four-test FDR family. Adjusted group effects were expressed as intermediate minus amyloid-positive; standardized effects used for cross-analysis comparison were obtained by dividing the group coefficient and its SE by the corresponding model residual SD.

#### 2.6.2 Descriptive regional distribution of cortical deviations

Before region-wise inferential testing, the spatial distribution of cortical normative deviations was characterized descriptively within the amyloid-negative, intermediate, and amyloid-positive groups. For each of the 204 cortical molecular-system features, the group-wise mean *Z*^std^ was calculated to retain the direction of the average deviation. In parallel, regional extreme-deviation prevalence was defined as the percentage of participants within each amyloid-burden stratum with an absolute locally standardized deviation exceeding the conventional |*Z*| > 1.96 threshold on the locally standardized scale. For each feature and group, the number of participants exceeding this threshold and the corresponding prevalence were tabulated. Intermediate minus amyloid-positive prevalence differences were additionally calculated in percentage points as descriptive contrasts; no region-wise inferential test or multiplicity correction was applied to these prevalence differences.

#### 2.6.3 Regional deviations from the locally calibrated normative reference

Region-wise reference-relative analyses were performed using two complementary deviation-score representations.

First, because *Z*^std^ was centered and scaled feature-by-feature and batch-by-batch using the local CDR = 0, CL < 10 reference participants, the reference location for each feature was zero by construction. For each of the 204 cortical features, separate two-sided one-sample *t*-tests evaluated whether the mean *Z*^std^ differed from zero in the intermediate-burden group and in the amyloid-positive group. FDR correction was applied separately across the fixed 204-feature family for each amyloid group. These tests quantified deviation from the locally calibrated amyloid-negative group. Because the reference location was estimated from the local reference participants, these one-sample tests do not propagate the sampling uncertainty of that estimate and were interpreted as descriptive, reference-anchored contrasts.

#### 2.6.4 Regional differences between intermediate amyloid burden and amyloid positivity

The predefined independent regional comparison directly contrasted participants with 10 ≤ CL < 30 and those with CL ≥ 30. For each of the 204 cortical features, the primary model used *Z*^std^ as the dependent variable and included amyloid group, age, sex, and acquisition batch as predictors. The reported group effect was the HC3-adjusted intermediate minus amyloid-positive difference with robust SE, 95% confidence interval (CI), and two-sided *p*-value. Group-effect *p*-values were FDR corrected across the fixed 204-feature family. Standardized group effects were obtained by dividing the group coefficient and its SE by the model residual SD.

To assess residual downstream covariate contributions, age and sex were evaluated from their HC3 coefficient tests, whereas acquisition batch, represented by multiple indicator variables, was assessed using a robust joint Wald test. Age, sex, and batch-effect *p*-values were each FDR corrected across the 204 regional models for descriptive multiplicity control. These diagnostics were not used as a data-driven criterion for retaining or removing covariates.

The regional analysis was then repeated using a group-only HC3 model for each feature. Adjusted and unadjusted group coefficients, CIs, and FDR conclusions were compared directly.

#### 2.6.5 Associations with continuous Centiloid burden and with APOE *ɛ*4

Continuous amyloid analyses were restricted to cognitively unimpaired participants with CL ≥ 10, thereby excluding the amyloid-negative transfer-reference group. For each of the 204 cortical features, *Z*^std^ was regressed on CL burden, age, sex, and acquisition batch. CL was scaled so that coefficients represented the expected change in normative deviation per 10-CL increase. Regional *p*-values were FDR corrected across the fixed 204-feature family. Continuous associations were also evaluated for global, DAT, NET, and SERT RADB and extreme-deviation burden, with FDR correction across the four scopes separately for each outcome.

APOE *ɛ*4 carrier status was examined as a modifier of the association between amyloid burden and normative deviation. In cognitively unimpaired participants with CL ≥ 10, RADB was modeled as a function of CL burden, APOE *ɛ*4 carrier status, their interaction, age, sex, and acquisition batch. The interaction term was the effect of interest. Tests were performed separately for the global, DAT, NET, and SERT scopes, with FDR correction across the four interaction tests.

Additional sensitivity analyses addressing diagnostic feature filtering, deviation-score representation, downstream covariate specification, and RADB distributional handling are reported in Supplementary Section A.2.

Statistical analyses were implemented in Python using numpy, pandas, scipy, statsmodels, and matplotlib. Cortical surface visualizations used the Enhancing NeuroImaging Genetics through Meta-Analysis (ENIGMA) Toolbox (Larivière et al., 2021).

## 3 Results

### 3.1 Healthy normative reference cohort and internal model validation

The final healthy normative reference cohort comprised 4,152 participants from seven datasets, including 3,324 participants in the training set and 828 in the held-out healthy test set (Table 6). The healthy reference sample spanned 18.0–89.8 years of age (mean 45.7 ± 18.7 years) and included 2,408 females and 1,744 males.

**Table 6:**
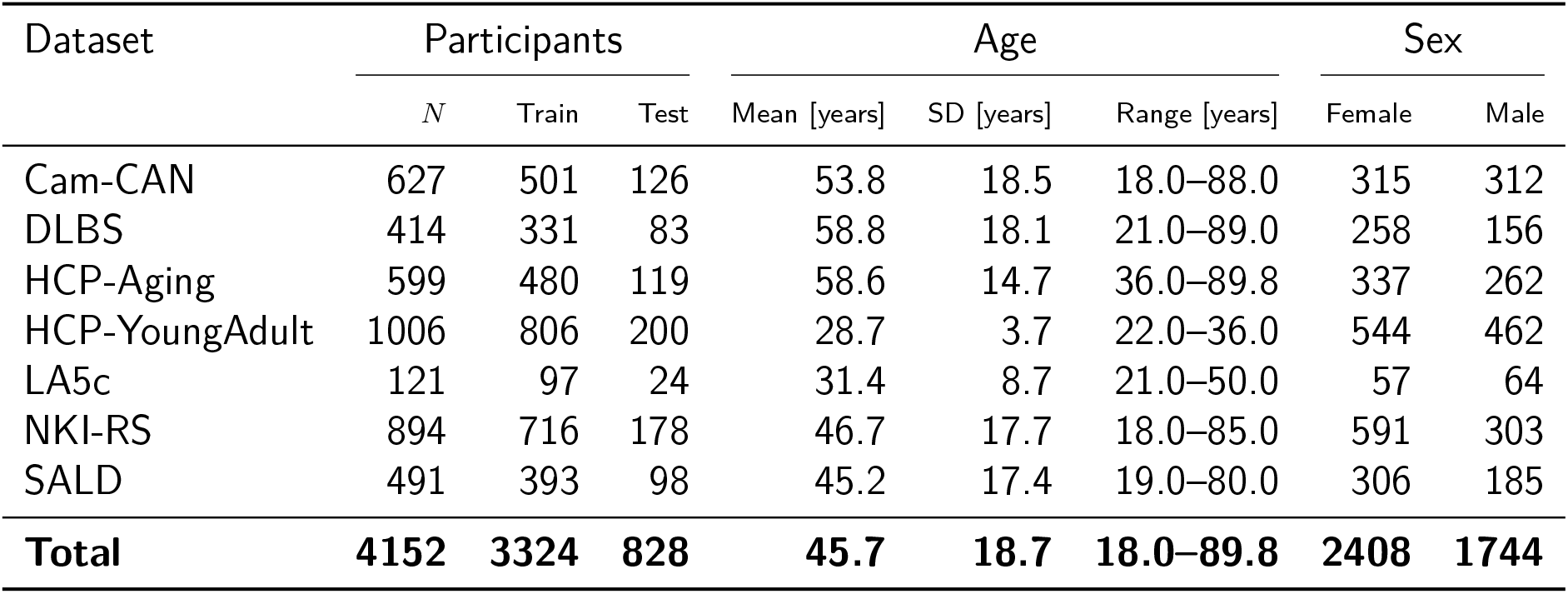
Healthy normative reference cohort characteristics by dataset.

| Dataset | Participants |  |  | Age |  |  | Sex |  |
| --- | --- | --- | --- | --- | --- | --- | --- | --- |
|  | *N* | Train | Test | Mean [years] | SD [years] | Range [years] | Female | Male |
| Cam-CAN | 627 | 501 | 126 | 53.8 | 18.5 | 18.0–88.0 | 315 | 312 |
| DLBS | 414 | 331 | 83 | 58.8 | 18.1 | 21.0–89.0 | 258 | 156 |
| HCP-Aging | 599 | 480 | 119 | 58.6 | 14.7 | 36.0–89.8 | 337 | 262 |
| HCP-YoungAdult | 1006 | 806 | 200 | 28.7 | 3.7 | 22.0–36.0 | 544 | 462 |
| LA5c | 121 | 97 | 24 | 31.4 | 8.7 | 21.0–50.0 | 57 | 64 |
| NKI-RS | 894 | 716 | 178 | 46.7 | 17.7 | 18.0–85.0 | 591 | 303 |
| SALD | 491 | 393 | 98 | 45.2 | 17.4 | 19.0–80.0 | 306 | 185 |
| **Total** | **4152** | **3324** | **828** | **45.7** | **18.7** | **18.0–89.8** | **2408** | **1744** |

All 204 predefined cortical NMs completed estimation. Of these, 199/204 (97.5%) satisfied the strict NM diagnostic criteria, including 65/68 DAT models, 67/68 NET models, and 67/68 SERT models (Table 7). The five models not meeting the strict diagnostic screen corresponded to DAT-enriched left entorhinal, left inferior temporal, and right entorhinal cortex, NET-enriched left lingual cortex, and SERT-enriched left temporal pole. All five models completed estimation but failed one or more of the predefined posterior-sampling or held-out calibration criteria. Held-out predictive performance varied across cortical features and molecular systems (Figure 1). Across all 204 models, the median EXPV was 0.174 (interquartile range (IQR) 0.048–0.336), the median SMSE was 0.827 (IQR 0.665–0.953), and the median MSLL was −4.497 (IQR −4.708 to −4.347). Median EXPV was 0.210 (IQR 0.070–0.392) for DAT, 0.102 (IQR 0.006–0.261) for NET, and 0.175 (IQR 0.056–0.346) for SERT. Held-out deviation-score calibration was close to the expected standardized distribution across models: the median test-set mean Z-score was 0.003 (IQR −0.019 to 0.030), the median Z-score SD was 1.006 (IQR 0.987–1.033), and the median maximum absolute Z-score was 4.60 (IQR 4.16–5.83).

**Figure 1:**
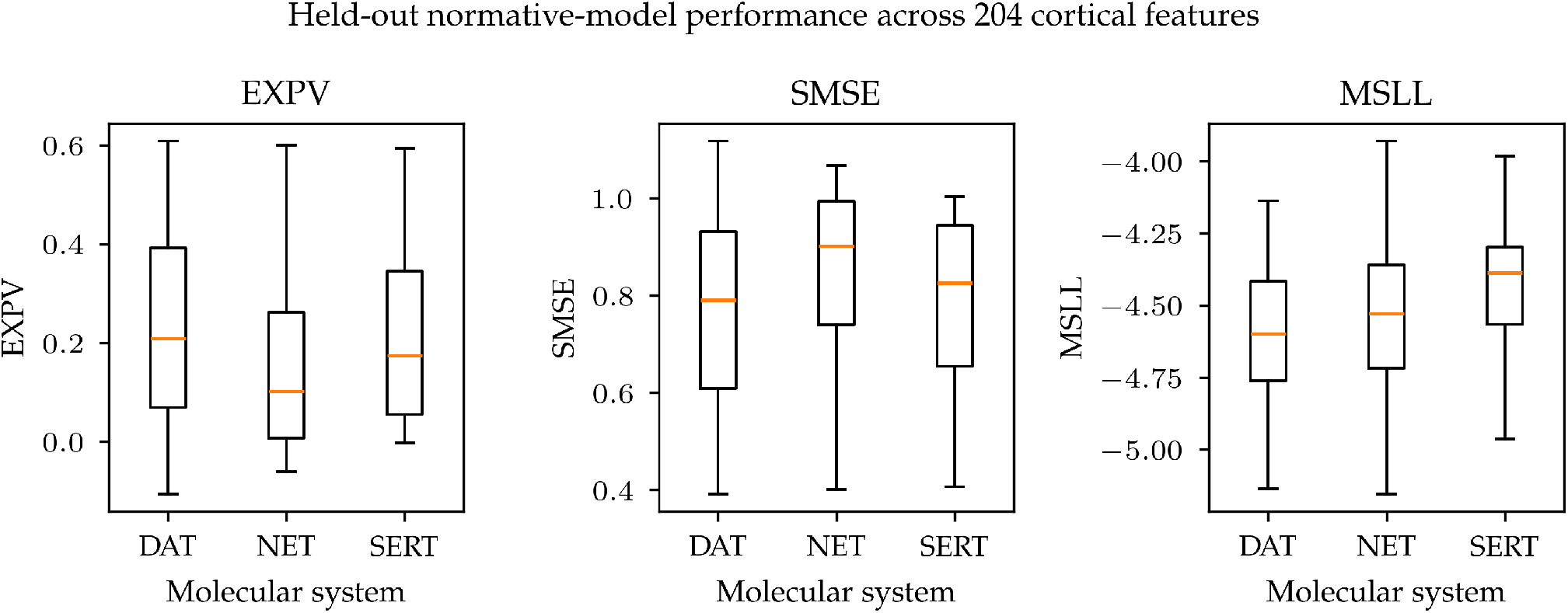
Held-out normative-model performance across the 204 cortical molecular-enriched functional-connectivity features. Boxplots summarize EXPV, SMSE, and MSLL across the 68 DAT-, 68 NET-, and 68 SERT-enriched cortical models.

**Table 7:** Held-out validation and strict diagnostic-screen summary for the 204 cortical normative models.

| System | Models | QC pass |  | EXPV | SMSE | MSLL | Test-Z SD |
| --- | --- | --- | --- | --- | --- | --- | --- |
|  |  | *n* | % | median [IQR] | median [IQR] | median [IQR] | median [IQR] |
| DAT | 68 | 65 | 95.6 | 0.210 [0.070, 0.392] | 0.791 [0.609, 0.932] | −4.600 [−4.762, −4.416] | 1.039 [1.011, 1.065] |
| NET | 68 | 67 | 98.5 | 0.102 [0.006, 0.261] | 0.902 [0.739, 0.994] | −4.528 [−4.718, −4.360] | 0.990 [0.970, 1.006] |
| SERT | 68 | 67 | 98.5 | 0.175 [0.056, 0.346] | 0.825 [0.654, 0.944] | −4.387 [−4.566, −4.298] | 1.004 [0.986, 1.022] |
| All | 204 | 199 | 97.5 | 0.174 [0.048, 0.336] | 0.827 [0.665, 0.953] | −4.497 [−4.708, −4.347] | 1.006 [0.987, 1.033] |

QC pass denotes satisfaction of the predefined posterior/sampler and held-out deviation-score diagnostic criteria described in Section 2.5.2. Performance summaries include all 204 completed models, irrespective of strict QC status. IQR = interquartile range; EXPV = explained variance; SMSE = standardized mean squared error; MSLL = mean standardized log loss.

### 3.2 AMYPAD-PNHS analytical cohort and transfer diagnostics

The downstream cortical analysis included 529 unique AMYPAD-PNHS cognitively unimpaired participants. Of these, 352 had CL < 10, 111 had 10 ≤ CL < 30, and 66 had CL ≥ 30 (Table 8). Mean age was 64.3 ± 7.2, 66.0 ± 6.6, and 69.4 ± 6.0 years in the amyloid-negative, intermediate-amyloid-burden, and amyloid-positive groups, respectively. Median CL values were 0.78 (IQR −3.73 to 4.67), 14.95 (IQR 12.37–20.04), and 54.07 (IQR 40.71–67.76), respectively. Mean Mini-Mental State Examination (MMSE) scores were 29.13, 29.30, and 28.92, respectively. Across the 529 participants in the final analytical cohort, the median absolute interval between MRI and amyloid PET acquisition was 37 days (IQR 5–106; range 0–357 days). All three amyloid-burden strata were represented in each acquisition batch (Table 8).

**Table 8:** Characteristics of the cognitively unimpaired AMYPAD-PNHS analytical sample and its distribution across acquisition batches.

| **A. Participant characteristics by amyloid group** |  |  |  |  |  |  |
| --- | --- | --- | --- | --- | --- | --- |
| Amyloid group | *N* | Age | Sex | CL | MMSE | Education |
|  |  | mean (SD) [years] | female / male | median [IQR] | mean (SD) [*n*] | mean (SD) |
| CL < 10 | 352 | 64.3 (7.2) | 213 / 139 | 0.78 [−3.73, 4.67] | 29.13 (1.01) [344] | 14.87 (4.28) |
| 10 ≤ CL < 30 | 111 | 66.0 (6.6) | 61 / 50 | 14.95 [12.37, 20.04] | 29.30 (0.93) [111] | 14.95 (4.04) |
| CL ≥ 30 | 66 | 69.4 (6.0) | 40 / 26 | 54.07 [40.71, 67.76] | 28.92 (1.23) [65] | 14.23 (4.47) |

| **B. Participant counts by acquisition batch** |  |  |  |  |
| --- | --- | --- | --- | --- |
| Acquisition batch | CL < 10 | 10 ≤ CL < 30 | CL ≥ 30 | Total |
| BBRC-ALFA | 138 | 48 | 24 | 210 |
| FACEHBI | 148 | 21 | 18 | 187 |
| VUmc | 66 | 42 | 24 | 132 |

### 3.2 AMYPAD-PNHS cohort and transfer diagnostics

All 612 cortical feature–batch transfer tasks (204 cortical features × 3 AMYPAD acquisition batches) completed successfully. Of these, 541/612 (88.4%) satisfied the complete predefined transfer-QC criteria, which jointly required acceptable posterior and sampler diagnostics and adequate calibration of locally standardized deviation scores in held-out local-reference participants (Table 9). The remaining 71/612 transfers failed at least one component of the strict transfer-QC screen.

**Table 9:** AMYPAD transfer quality-control summary across molecular systems and acquisition batches.

| Acquisition batch | DAT |  | NET |  | SERT |  | Overall |  |
| --- | --- | --- | --- | --- | --- | --- | --- | --- |
|  | *n/N* | % | *n/N* | % | *n/N* | % | *n/N* | % |
| BBRC-ALFA | 61/68 | 89.7 | 65/68 | 95.6 | 66/68 | 97.1 | 192/204 | 94.1 |
| FACEHBI | 51/68 | 75.0 | 59/68 | 86.8 | 62/68 | 91.2 | 172/204 | 84.3 |
| VUmc | 54/68 | 79.4 | 62/68 | 91.2 | 61/68 | 89.7 | 177/204 | 86.8 |
| All batches | 166/204 | 81.4 | 186/204 | 91.2 | 189/204 | 92.6 | 541/612 | 88.4 |

Transfer-QC performance varied across acquisition batches and molecular systems. Aggregated across the three molecular systems, 192/204 transfers (94.1%) satisfied transfer QC in BBRC-ALFA, 172/204 (84.3%) in FACEHBI, and 177/204 (86.8%) in VUmc. Aggregated across acquisition batches, the corresponding pass fractions were 166/204 (81.4%) for DAT, 186/204 (91.2%) for NET, and 189/204 (92.6%) for SERT. At the individual system-by-batch level, transfer-QC pass fractions ranged from 75.0% for DAT in FACEHBI to 97.1% for SERT in BBRC-ALFA (Table 9).

When normative-model and transfer diagnostics were considered jointly, 535/612 feature–batch pairs satisfied both screens. Requiring a feature to satisfy the complete normative-model and transfer-diagnostic criteria in every AMYPAD batch yielded a strict cross-batch diagnostic sensitivity domain of 157 cortical features, comprising 48 DAT, 55 NET, and 54 SERT features. Consistent with the atlas-complete primary analysis strategy, diagnostic-pass status was treated as information about model reliability rather than as an anatomical feature-selection rule. The primary analyses therefore retained the 204 cortical features with finite transferred deviation scores, whereas analyses restricted to the 157-feature diagnostic domain were used to assess sensitivity to the strict diagnostic thresholds (see Supplementary Section A.2). Complete finite profiles across all 204 features were available for all 529 participants included in the primary subject-level analyses.

### 3.3 Subject-level deviation burden across amyloid groups

We first examined whether the multiregional cortical deviation profile differed across amyloid-burden groups when summarized at the subject level. Across the three amyloid groups, the Welch omnibus test identified a difference in DAT-specific RADB (*F* = 5.49, *p* = 0.005, *q* = 0.020), whereas the global, NET, and SERT omnibus tests did not survive FDR correction (global *q*= 0.113, NET *q*= 0.364, SERT *q*= 0.151). DAT RADB means were 1.309 in the amyloid-negative group, 1.296 in the intermediate-burden group, and 1.121 in the amyloid-positive group. In pairwise analyses, DAT RADB was lower in the amyloid-positive than in the amyloid-negative reference group (mean difference = −0.188, 95% CI [−0.302, −0.074], *p* = 0.001, *q* = 0.006), and global RADB was also lower in the amyloid-positive than in the amyloid-negative group (difference = −0.150, 95% CI [−0.280, −0.019], *p* = 0.025, *q* = 0.050). No intermediate amyloid burden versus amyloid-negative RADB comparison survived FDR correction, and no unadjusted intermediate versus amyloid-positive pairwise comparison survived FDR correction across the four scopes (minimum *q* = 0.072). The omnibus and amyloid-positive versus amyloid-negative pairwise results are summarized in Table 10, and the unadjusted group distributions are shown in Figure 2.

**Figure 2:**
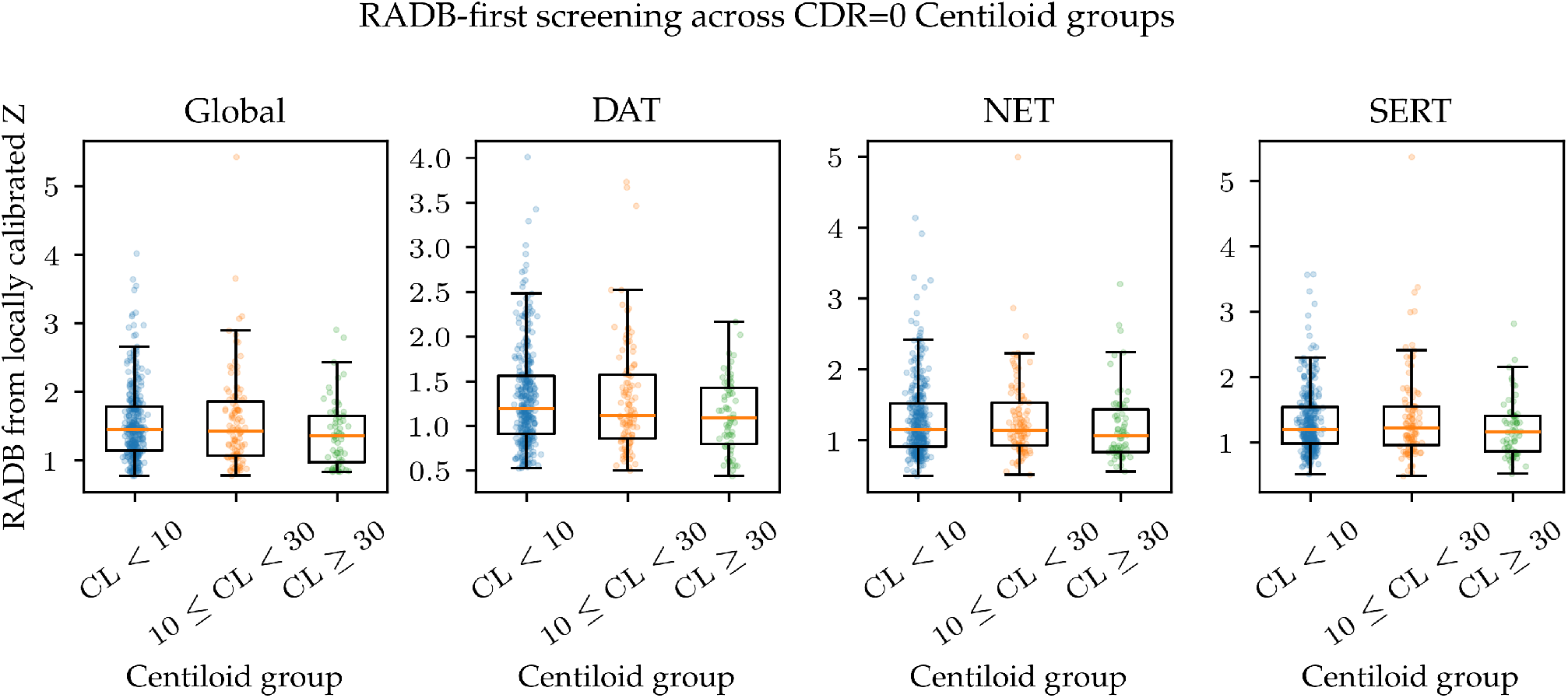
Ranked absolute deviation burden (RADB) across the three amyloid groups for the global cortical domain and DAT-, NET-, and SERT-specific feature subsets. Points represent individual participants and box-plots summarize the unadjusted distributions.

**Table 10:** RADB-first screening across the three amyloid groups.

| Scope | Omnibus Welch test | | | CL $\geq 30$ minus CL $< 10$ | | |
| --- | --- | --- | --- | --- | --- | --- |
| | $F$ | $p$ | FDR $q$ | Difference | 95% CI | FDR $q$ |
| Global | 2.929 | 0.057 | 0.113 | −0.150 | [−0.280, −0.019] | 0.050 |
| DAT | 5.485 | 0.005 | **0.020** | −0.188 | [−0.302, −0.074] | **0.006** |
| NET | 1.017 | 0.364 | 0.364 | −0.098 | [−0.237, 0.040] | 0.161 |
| SERT | 2.213 | 0.113 | 0.151 | −0.106 | [−0.223, 0.012] | 0.104 |

Omnibus tests are heteroscedasticity-robust Welch one-way comparisons across CL $< 10$ , $10 \leq \text{CL} < 30$ , and CL $\geq 30$ . Pairwise values shown in the final three columns correspond to CL $\geq 30$ minus the local CL $< 10$ transfer reference group. FDR correction was applied across the four scopes separately for the omnibus family and for each pairwise contrast.

The predefined independent intermediate versus amyloid-positive comparison was additionally evaluated with HC3 regression adjusted for age, sex, and acquisition batch (Table 11). RADB was higher in the intermediate group across all four scopes, but none of the adjusted RADB contrasts survived FDR correction: global difference = 0.182 (95% CI 0.016–0.349, *p* = 0.031, *q* = 0.062), DAT = 0.165 (95% CI 0.015–0.315, *p* = 0.031, *q* = 0.062), NET = 0.095 (95% CI −0.067–0.258, *p* = 0.250, *q* = 0.250), and SERT = 0.169 (95% CI 0.003–0.335, *p* = 0.047, *q*= 0.062). For extreme-deviation burden, DAT showed a greater burden in the intermediate than in the amyloid-positive group (adjusted difference = 0.032, 95% CI 0.007–0.057, *p* = 0.011, *q* = 0.044), corresponding to approximately 3.2 percentage points, or about two of the 68 DAT cortical features. The global and SERT estimates were also positive but did not survive FDR correction (global difference = 0.023, *q* = 0.072; SERT difference = 0.034, *q* = 0.072), whereas the NET difference was 0.003 (*q* = 0.837).

**Table 11:**
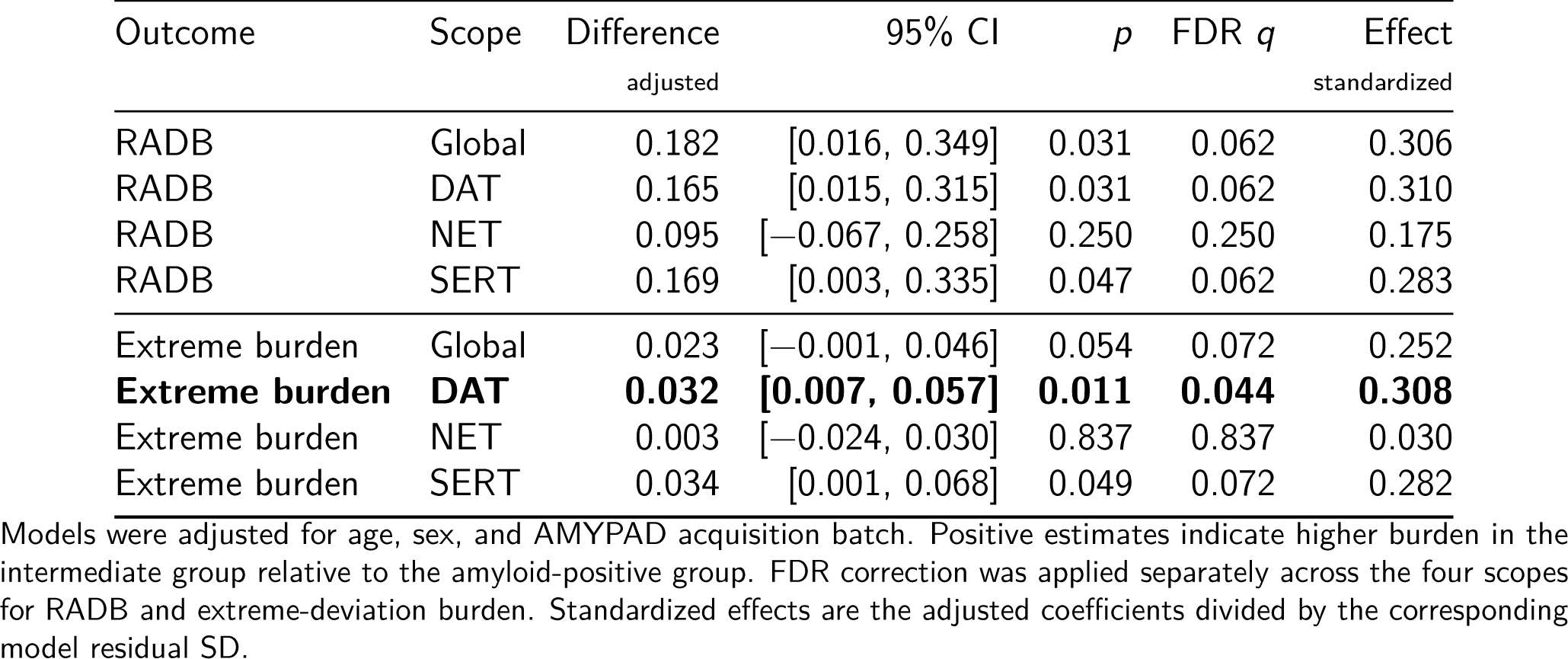
Adjusted subject-level deviation-burden comparison between intermediate and amyloid-positive.

### 3.4 Spatial distribution of locally standardized cortical deviations

The spatial distributions of locally standardized cortical deviation scores across the three amyloid groups are shown in Figures 3 and 4. Figure 3 displays, for each DAT-, NET-, and SERT-enriched cortical feature, the group-wise mean *Z*^std^, thereby retaining the direction of the average deviation from the locally calibrated normative reference. Figure 4 displays the complementary regional prevalence of extreme deviations, defined as the percentage of participants within each group with |*Z*^std^| > 1.96. The amyloid-negative group is shown descriptively because these participants contributed to local transfer and calibration.

**Figure 3:**
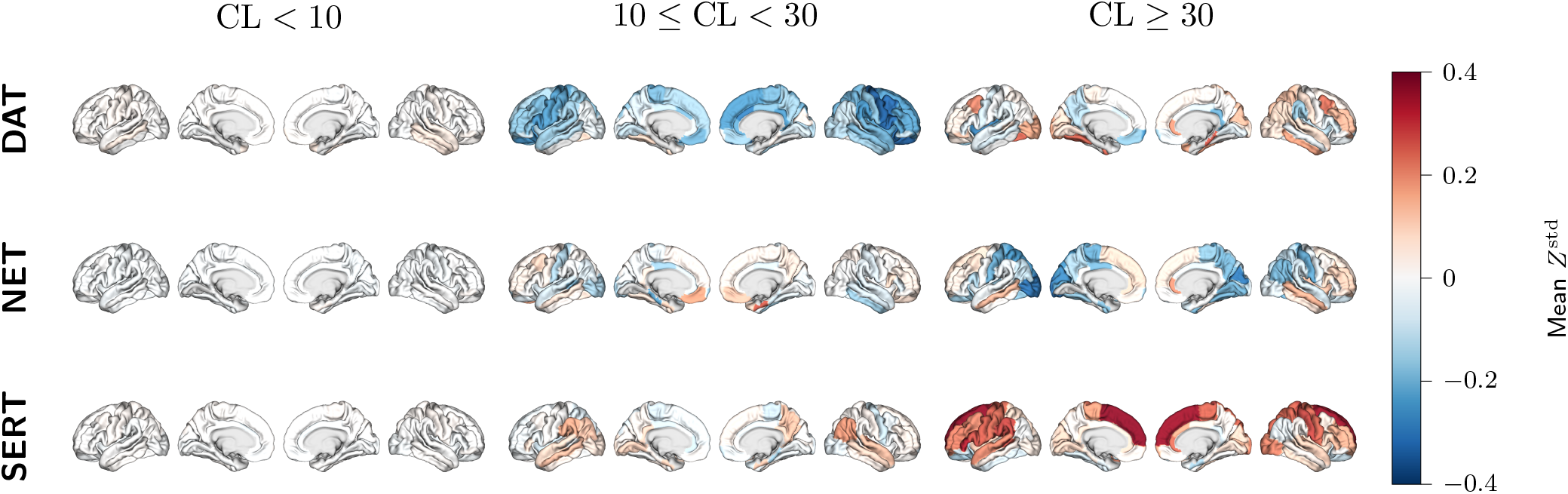
Mean locally standardized cortical normative deviations across the three Centiloid groups. Rows correspond to DAT-, NET-, and SERT-enriched cortical features, and columns correspond to the amyloid-negative, intermediate-amyloid-burden, and amyloid-positive groups, respectively. For each cortical region, values represent the group-wise mean *Z*^std^. Positive values indicate mean deviations above the locally calibrated normative reference, whereas negative values indicate mean deviations below it.

**Figure 4:**
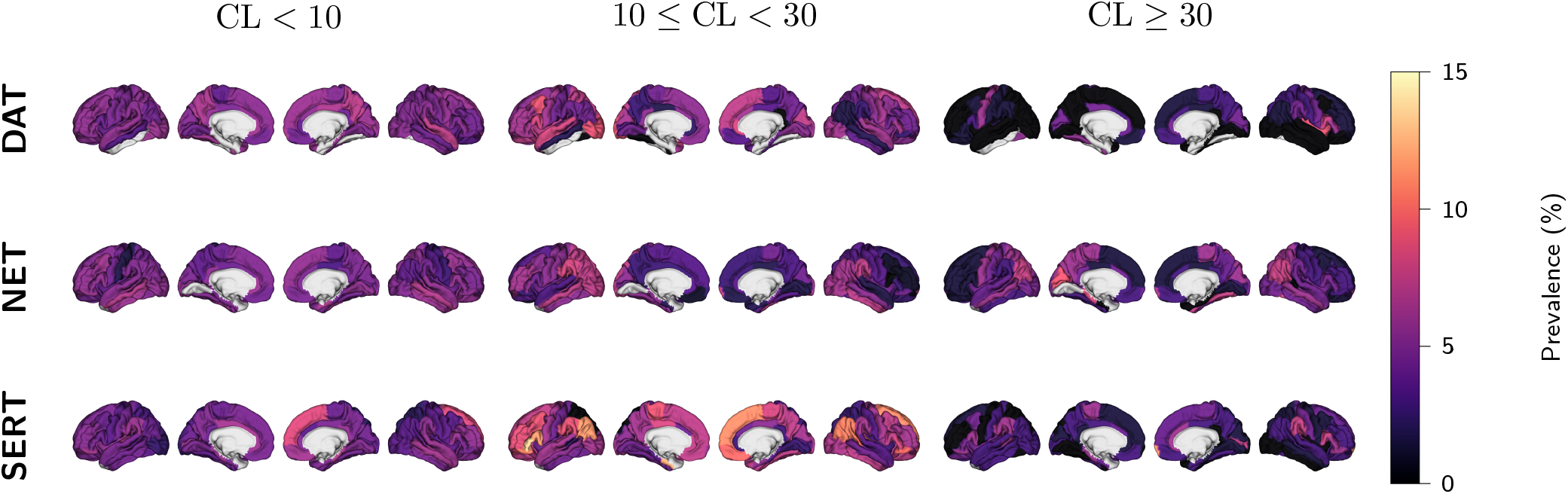
Prevalence of extreme locally standardized cortical normative deviations across the three amyloid groups. Rows correspond to DAT-, NET-, and SERT-enriched cortical features, and columns correspond to the amyloid-negative, intermediate-amyloid-burden, and amyloid-positive groups, respectively. For each cortical region, values represent the percentage of participants with |*Z*^std^| > 1.96. The prevalence measure is unsigned and therefore reflects the frequency of deviations exceeding the predefined threshold irrespective of whether the deviation was positive or negative.

The unthresholded mean maps showed spatially heterogeneous and molecular-system-specific patterns. Among the two target amyloid groups, mean *Z*^std^ values across the 204 cortical features ranged from −0.354 to 0.193 in the intermediate-burden group and from −0.293 to 0.356 in the amyloid-positive group. For DAT-enriched connectivity, the most negative intermediate-group mean was observed in right caudal middle frontal cortex (−0.354); the same intermediate group also showed negative reference-relative deviations in right pars opercularis, lateral orbitofrontal, and precentral cortices in the subsequent inferential analysis. For NET-enriched connectivity, right banks of the superior temporal sulcus showed a negative intermediate-group mean of −0.340, whereas right cuneus was the most negative feature in the amyloid-positive group (−0.293). For SERT-enriched connectivity, the most positive intermediate group mean was observed in right transverse temporal cortex (0.193), while left superior frontal cortex was the most positive feature in the amyloid-positive group (0.356).

Extreme-deviation prevalence was likewise regionally heterogeneous. Across all 204 features, prevalence ranged from 1.7% to 7.1% in the CL < 10 local-reference group, from 0.9% to 13.5% in the intermediate-burden group, and from 0% to 9.1% in the amyloid-positive group. In the local-reference group, the maximum prevalence was 7.1%, observed in DAT-enriched right precuneus and SERT-enriched left transverse temporal cortex (25/352 participants each). In the intermediate-burden group, the maximum was 13.5% in SERT-enriched left pars triangularis (15/111). In the amyloid-positive group, the maximum was 9.1%, shared by DAT-enriched right insula, NET-enriched left cuneus, and NET-enriched left pericalcarine cortex (6/66 each). The largest descriptive intermediate-burden minus amyloid-positive prevalence differences were +9.0% in DAT-enriched left lateral occipital cortex and SERT-enriched left rostral middle frontal cortex; the largest difference in the opposite direction was −4.6% in NET-enriched left pericalcarine cortex.

### 3.5 Regional deviations from the locally calibrated normative reference

Region-wise reference-relative tests were performed separately in the intermediate and amyloid-positive groups. Using *Z*^std^, five of the 204 cortical molecular-system features survived FDR correction in the intermediate group (Table 12; Figure 5). All five effects were negative. Four involved DAT-enriched connectivity in the right hemisphere — caudal middle frontal (mean *Z* = −0.354, 95% CI [−0.534, −0.174], *p* < 0.001, *q* = 0.027), pars opercularis (mean *Z* = −0.313, 95% CI [−0.486, −0.140], *p* < 0.001, *q* = 0.027), lateral orbitofrontal (mean *Z* = −0.327, 95% CI [−0.510, −0.144], *p*< 0.001, *q* = 0.027), and precentral cortex (mean *Z* = −0.336, 95% CI [−0.526, −0.146], *p* < 0.001, *q* = 0.027) — and one involved NET-enriched connectivity in the right banks of the superior temporal sulcus (mean *Z* = −0.340, 95% CI [−0.529, −0.150], *p* < 0.001, *q* = 0.027). In the amyloid-positive group, one feature survived correction: SERT-enriched left superior frontal cortex showed a positive mean deviation (mean *Z* = 0.356, 95% CI 0.174–0.539, *p* < 0.001, *q* = 0.047).

**Figure 5:**
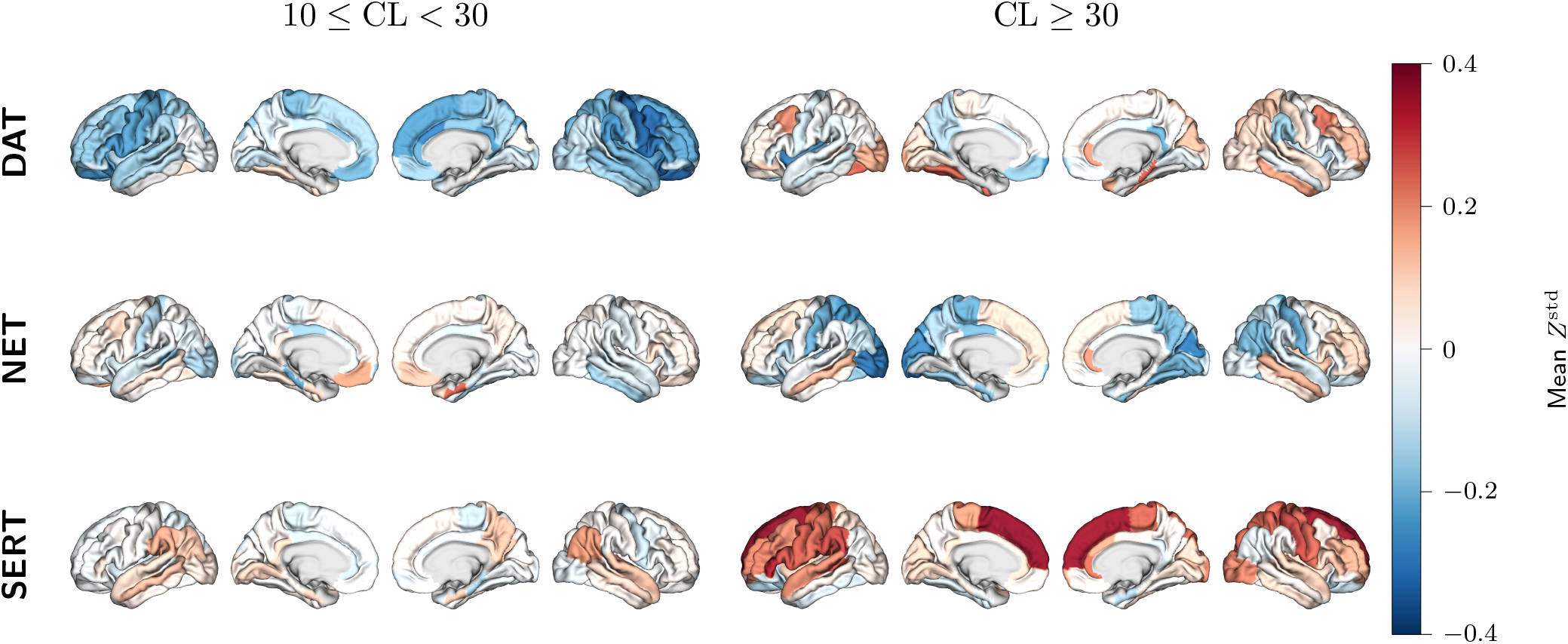
Unthresholded cortical maps of mean locally standardized normative deviation relative to the local amyloid-negative reference. The left column shows participants with intermediate amyloid burden and the right column amyloid-positive participants. Values are mean *Z*^std^; negative values indicate deviations below and positive values deviations above the local reference location. Color scales are centered on zero and generated separately for each map, so color intensity should not be compared quantitatively across panels. FDR-significant regions are reported in Table 12.

**Table 12:** FDR-corrected regional deviations.

| Amyloid group | System | Hemisphere | Region | Mean Z | 95% CI | *t* | *p* | FDR *q* |
| --- | --- | --- | --- | --- | --- | --- | --- | --- |
| $10 \leq \text{CL} < 30$ | DAT | RH | Caudal middle frontal | -0.354 | [-0.534, -0.174] | -3.893 | <0.001 | 0.027 |
| $10 \leq \text{CL} < 30$ | DAT | RH | Pars opercularis | -0.313 | [-0.486, -0.140] | -3.586 | <0.001 | 0.027 |
| $10 \leq \text{CL} < 30$ | NET | RH | Banks of superior temporal sulcus | -0.340 | [-0.529, -0.150] | -3.554 | <0.001 | 0.027 |
| $10 \leq \text{CL} < 30$ | DAT | RH | Lateral orbitofrontal | -0.327 | [-0.510, -0.144] | -3.548 | <0.001 | 0.027 |
| $10 \leq \text{CL} < 30$ | DAT | RH | Precentral | -0.336 | [-0.526, -0.146] | -3.507 | <0.001 | 0.027 |
| $\text{CL} \geq 30$ | SERT | LH | Superior frontal | 0.356 | [0.174, 0.539] | 3.903 | <0.001 | 0.047 |

Tests were two-sided one-sample *t*-tests against zero, performed separately within the intermediate and amyloid-positive groups. FDR correction used a fixed 204-feature family separately for each group.

### 3.6 Regional differences between intermediate amyloid burden and amyloid-positive groups

The independent regional comparison comprised 177 participants who did not contribute to the local CL < 10 transfer-reference group: 111 with 10 ≤ CL < 30 and 66 with CL ≥ 30. The primary HC3 analysis adjusted for age, sex, and acquisition batch identified a group difference in DAT-enriched right caudal middle frontal cortex that survived FDR correction (Table 13; Figure 6). Locally standardized deviation scores were lower in the intermediate than in the amyloid-positive group (adjusted intermediate-minus-positive difference = −0.582, HC3 SE = 0.147, 95% CI [−0.871, −0.293], *p* < 0.001, *q* = 0.016; standardized effect = −0.635). Sixteen regional models had nominal *p* < 0.05 before correction (8 DAT, 1 NET, and 7 SERT); the right caudal middle frontal DAT feature was the effect that met the corrected regional significance threshold.

**Figure 6:**
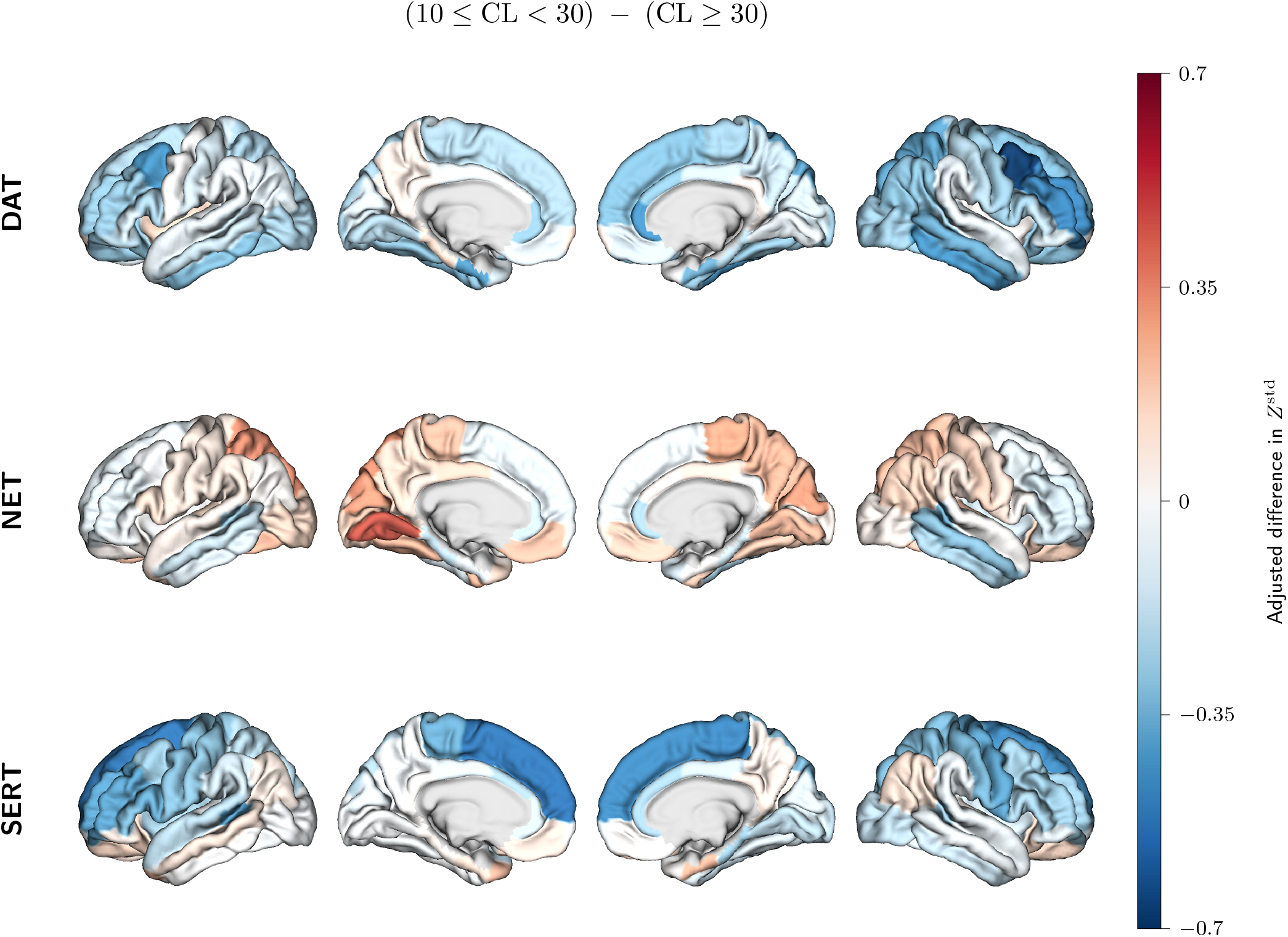
Unthresholded cortical maps of the adjusted intermediate minus amyloid-positive group effect for DAT-, NET-, and SERT-enriched locally standardized normative deviations. Models included age, sex, and acquisition batch. Negative values indicate lower deviation scores in the intermediate group. Only DAT-enriched right caudal middle frontal cortex survived FDR correction across the 204-feature family.

**Table 13:**
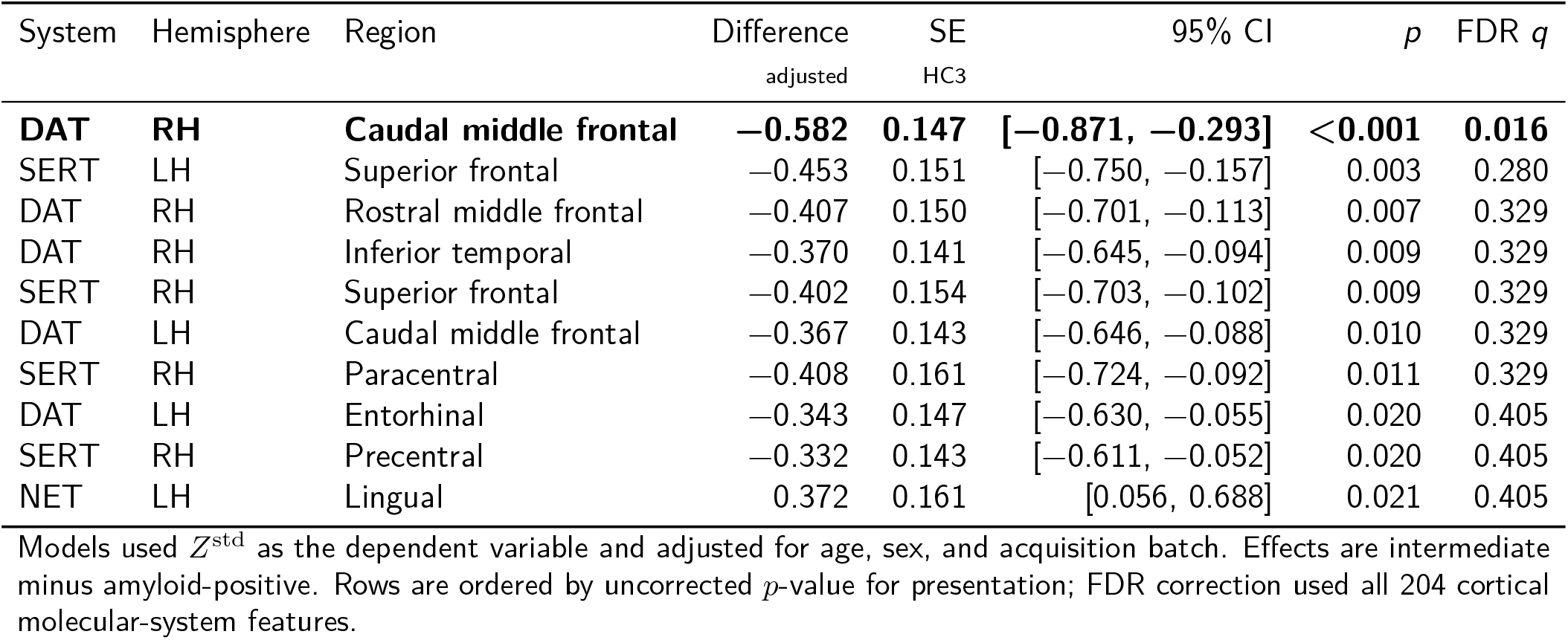
Cortical features with the strongest adjusted differences between intermediate and amyloid-positive groups.

In the covariate tests from the 204 adjusted regional models, age was nominally associated with 25 features and remained significant after FDR correction for two: DAT-enriched left entorhinal cortex and SERT-enriched right rostral middle frontal cortex (both *q* = 0.030). Sex was nominally associated with 25 features, with none surviving FDR correction. The joint acquisition-batch effect was nominally significant in 21 features and remained significant after FDR correction for NET-enriched right rostral middle frontal cortex (*q* < 0.001). For the DAT-enriched right caudal middle frontal feature with the statistically significant group effect, the age, sex, and joint batch terms had *p* = 0.317, 0.187, and 0.668, respectively.

In the group-only HC3 analysis, DAT-enriched right caudal middle frontal cortex remained the only significant feature after FDR correction (intermediate-minus-positive difference = −0.539, HC3 SE = 0.139, 95% CI [−0.811, −0.268], *p* < 0.001, *q* = 0.020). Across all 204 features, adjusted and unadjusted group coefficients were correlated at *r* = 0.947, 185/204 (90.7%) retained the same effect direction, and the median absolute coefficient change was 0.036. The correspondence between adjusted and group-only coefficients is shown in Supplementary Figure 8.

### 3.7 Associations with continuous Centiloid burden

Continuous CL analyses were restricted to the 177 cognitively unimpaired participants with CL ≥ 10 and therefore tested linear associations within the intermediate-to-amyloid-positive range rather than across the full CL distribution. No regional linear association survived FDR correction across the 204 cortical features (Table 14; Figure 7). The strongest regional association was observed for DAT-enriched right caudal middle frontal cortex, where the locally calibrated deviation score increased by 0.101 units per 10-CL increment (HC3 SE = 0.029, 95% CI 0.045–0.157, *p* < 0.001, *q* = 0.088). Eleven regional features had nominal *p*< 0.05 (5 DAT, 1 NET, and 5 SERT), with none meeting the FDR-corrected threshold.

**Figure 7:**
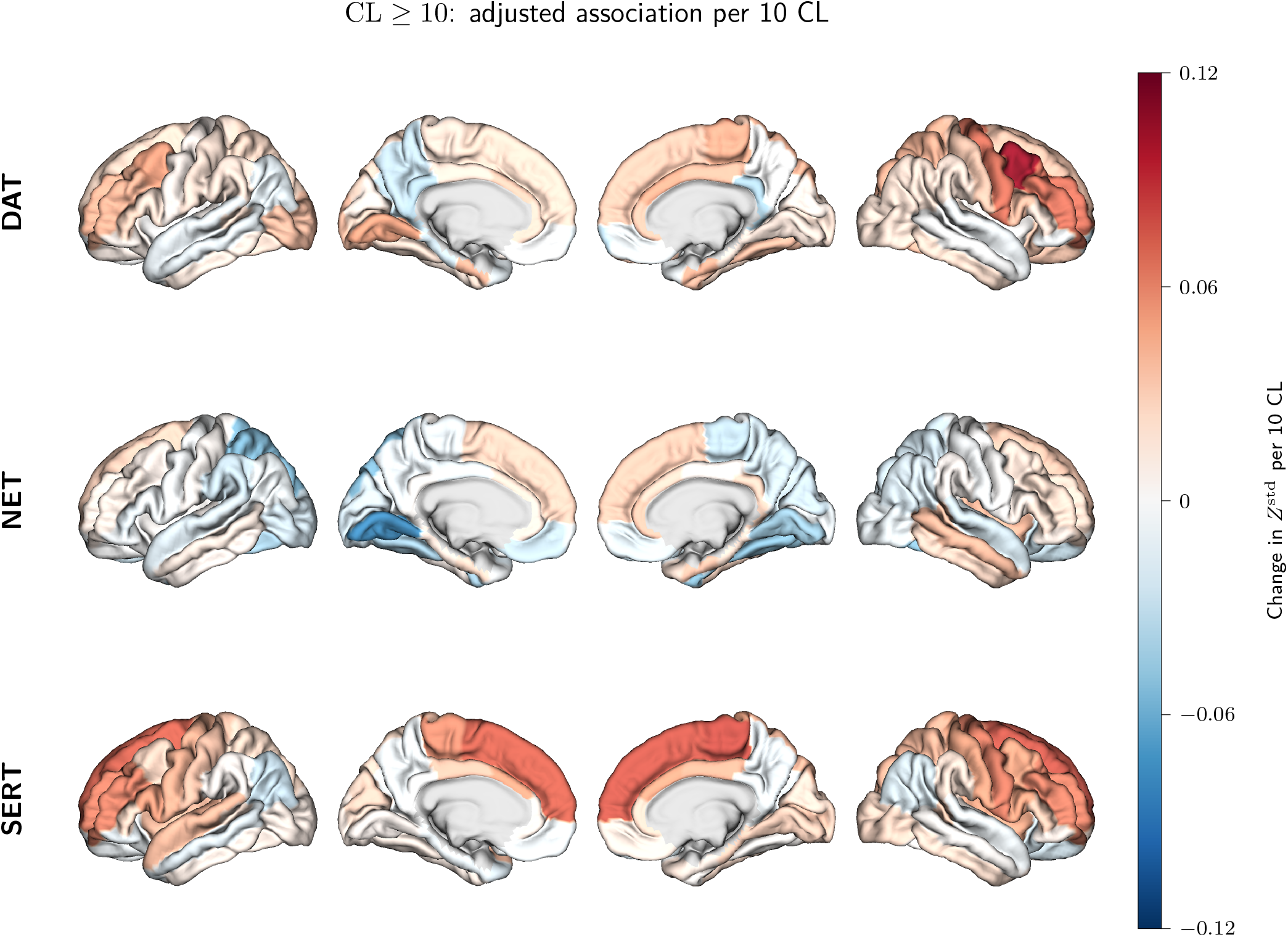
Unthresholded cortical maps of adjusted continuous Centiloid associations with DAT-, NET-, and SERT-enriched normative deviations among cognitively unimpaired participants with CL ≥ 10. Values represent the adjusted change in *Z*^std^ per 10-Centiloid increase after adjustment for age, sex, and acquisition batch. Color scales are centered on zero and are system-specific; therefore, color intensity should not be compared quantitatively across molecular systems. No regional association survived FDR correction across the 204-feature family.

**Table 14:** Cortical features with the strongest adjusted continuous Centiloid associations.

| System | Hemisphere | Region | Slope<br>per 10 CL | SE<br>HC3 | 95% CI | $p$ | FDR $q$ |
| --- | --- | --- | --- | --- | --- | --- | --- |
| DAT | RH | Caudal middle frontal | 0.101 | 0.029 | [0.045, 0.157] | <0.001 | 0.088 |
| SERT | RH | Paracentral | 0.077 | 0.028 | [0.023, 0.131] | 0.006 | 0.560 |
| SERT | RH | Precentral | 0.063 | 0.024 | [0.015, 0.111] | 0.010 | 0.634 |
| NET | LH | Lingual | -0.076 | 0.031 | [-0.138, -0.015] | 0.015 | 0.634 |
| DAT | RH | Rostral middle frontal | 0.066 | 0.028 | [0.010, 0.121] | 0.020 | 0.634 |
| SERT | RH | Superior frontal | 0.075 | 0.032 | [0.011, 0.139] | 0.021 | 0.634 |
| SERT | RH | Rostral middle frontal | 0.061 | 0.027 | [0.009, 0.113] | 0.022 | 0.634 |
| DAT | RH | Precentral | 0.066 | 0.031 | [0.006, 0.126] | 0.031 | 0.793 |
| SERT | LH | Paracentral | 0.056 | 0.027 | [0.003, 0.108] | 0.037 | 0.842 |
| DAT | LH | Lingual | 0.048 | 0.024 | [0.001, 0.096] | 0.045 | 0.850 |
| DAT | LH | Caudal middle frontal | 0.050 | 0.025 | [0.001, 0.100] | 0.046 | 0.850 |
| NET | LH | Superior parietal | -0.053 | 0.027 | [-0.106, 0.000] | 0.051 | 0.854 |
| NET | RH | Lingual | -0.043 | 0.022 | [-0.086, 0.001] | 0.056 | 0.854 |
| SERT | RH | Caudal middle frontal | 0.047 | 0.025 | [-0.002, 0.096] | 0.060 | 0.854 |
| SERT | RH | Posterior cingulate | 0.042 | 0.023 | [-0.003, 0.087] | 0.067 | 0.854 |

Models used $Z^{\text{std}}$ as the dependent variable and adjusted for age, sex, and acquisition batch. Slopes represent the expected change in deviation score per 10-CL increase among participants with $\text{CL} \geq 10$ . Rows are ordered by uncorrected $p$ -value for presentation only; FDR correction used the fixed family of all 204 cortical molecular-system features. No regional association survived at $q < 0.05$ .

Continuous CL burden was also not significantly associated with RADB or extreme-deviation burden after FDR correction (Table 15). The adjusted RADB slopes per 10 CL were −0.016 for the global scope (*q* = 0.393), −0.018 for DAT (*q* = 0.393), 0.002 for NET (*q* = 0.877), and −0.018 for SERT (*q* = 0.393). The corresponding extreme-deviation-burden slopes were −0.002 for global (*q* = 0.322), −0.004 for DAT (*q* = 0.226), 0.002 for NET (*q* = 0.519), and −0.005 for SERT (*q* = 0.247).

**Table 15:** Adjusted associations between continuous Centiloid burden and subject-level cortical deviation burden.

| Outcome | Scope | Slope<br>per 10 CL | 95% CI | $p$ | FDR $q$ |
| --- | --- | --- | --- | --- | --- |
| RADB | Global | −0.016 | [−0.046, 0.014] | 0.295 | 0.393 |
| RADB | DAT | −0.018 | [−0.043, 0.007] | 0.161 | 0.393 |
| RADB | NET | 0.002 | [−0.027, 0.031] | 0.877 | 0.877 |
| RADB | SERT | −0.018 | [−0.050, 0.013] | 0.253 | 0.393 |
| Extreme burden | Global | −0.002 | [−0.006, 0.002] | 0.242 | 0.322 |
| Extreme burden | DAT | −0.004 | [−0.008, 0.000] | 0.056 | 0.226 |
| Extreme burden | NET | 0.002 | [−0.003, 0.007] | 0.519 | 0.519 |
| Extreme burden | SERT | −0.005 | [−0.011, 0.001] | 0.124 | 0.247 |

Models included the 177 cognitively unimpaired participants with CL $\geq 10$ and were adjusted for age, sex, and acquisition batch. FDR correction was applied across the four scopes separately for RADB and extreme-deviation burden.

### 3.8 APOE *ɛ*4 interaction

The CL-by-APOE *ɛ*4 interaction was evaluated in 177 cognitively unimpaired participants with CL ≥ 10. None of the interaction terms survived correction across the global, DAT, NET, and SERT RADB scopes (all *q* ≥ 0.436). The largest interaction estimate was observed for DAT RADB (*β* = −0.044, 95% CI −0.098 to 0.010, *p* = 0.109, *q* = 0.436); the global, NET, and SERT interaction tests were also non-significant after FDR correction.

Sensitivity analyses of the principal categorical and continuous findings, including the strict cross-batch diagnostic domain and model-native *Z*^raw^ analyses, are reported in Supplementary Section A.2.1.

## 4 Discussion

This study addressed two complementary questions. First, we tested whether region-wise normative models of molecular-enriched FC could be estimated across heterogeneous healthy reference datasets and subsequently deployed in independently processed external data through local hierarchical adaptation. Second, we examined whether the transferred models could provide an informative individualized representation of functional variation during cognitively unimpaired stages of amyloid accumulation. The findings support the methodological feasibility of the framework and provide an initial biological application: most molecular-enriched phenotypes could be placed on a stable normative scale and transferred to previously unseen AMYPAD-PNHS acquisition environments, while the exploratory amyloid analyses identified modest, spatially and molecularly selective differences predominantly involving DAT-enriched FC. Together, these results support REACT-based NM as a distributed framework for individualized functional phenotyping, while indicating that the biological effects observed here require independent and longitudinal validation.

### 4.1 Normative reference modeling of molecular-enriched functional connectivity

The first step was to determine whether regional molecular-enriched functional phenotypes could be modeled reliably across the healthy adult lifespan. The high proportion of cortical models satisfying the predefined diagnostic criteria indicates that the HBR–SHASHb formulation was stable across a broad set of DAT-, NET-, and SERT-enriched phenotypes. The five models failing the strict diagnostic screen spanned the three molecular systems, but three involved DAT-enriched entorhinal and inferior temporal cortex, medial and inferior temporal regions in which gradient-echo BOLD signal is commonly reduced by susceptibility artifacts (Ojemann et al., 1997); these failures may therefore partly reflect regional signal quality rather than model specification.

Model adequacy in this setting should not be judged solely by the proportion of response variance explained by age and sex. The objective of NM is to estimate a calibrated conditional distribution from which individual deviations can be derived; substantial residual inter-individual variability is therefore expected, particularly for functional phenotypes (Maccioni et al., 2026; Marquand et al., 2016; Rutherford et al., 2022). The present held-out EXPV values are also consistent with the broader behavior of functional and molecular neuroimaging phenotypes. NMs of dopaminergic PET measures have explained only part of the between-subject variability using demographic covariates, and previous molecular-enriched functional NMs likewise show region- and system-dependent explained variance (Giacomel et al., 2025; Lawn et al., 2024). In this context, the close centering and scaling of the held-out deviation distributions is particularly relevant because systematic miscalibration would propagate directly into the individual abnormality scores used in the target cohort.

The four-parameter SHASHb further allowed conditional scale, skewness, and tail behavior to be represented rather than forcing all molecular-enriched phenotypes into a Gaussian conditional distribution (A. A. A. de Boer et al., 2024). This is important for individualized inference, where distributional misspecification could otherwise be interpreted as subject-level abnormality. The present models therefore extend our previous brain-age work from global age prediction to region-wise reference distributions that preserve anatomical location, molecular-system identity, direction, and extremeness of individual deviations (Pinamonti et al., 2026).

### 4.2 External transfer and local calibration

The first methodological objective of this study was to test whether the normative reference models could be transferred to previously unseen data. This represents a substantially different problem from internal held-out validation: a model distributed to a new hospital or medical imaging center may encounter scanner, acquisition, preprocessing, and site effects that were absent during reference-model estimation. The present design deliberately retained this challenge because AMYPAD-PNHS had been processed through an independently established upstream workflow rather than retrospectively routed through the complete normative-reference pipeline. Compatibility was imposed at the downstream phenotype level through common molecular templates, a common joint REACT design, the same cortical parcellation, and matched regional feature definitions.

Transfer was then performed using local amyloid-negative cognitively unimpaired reference participants to adapt the hierarchical models to each unseen acquisition batch. The high proportion of transfers satisfying the predefined diagnostics indicates that local HBR adaptation can absorb a substantial component of target-domain shift, although transferability was not uniform across features or batches. Transfer quality was lowest in FACEHBI (84.3% overall and 75.0% for DAT-enriched features), the only 1.5 T batch and the one with the longest repetition time (4 s) and fewest volumes, whereas all normative reference data were acquired at 3 T; the present design cannot separate field strength from other protocol and cohort differences, but this pattern suggests that target acquisitions far from the reference acquisition space are the most demanding for local adaptation. This distinction is important: the framework supports distributed deployment but should not be interpreted as zero-shot generalization. Local reference observations remain necessary to anchor the pretrained normative scale to the new acquisition environment, and explicit documentation of compatible downstream feature definitions and feature-level diagnostics are required to identify phenotypes for which the resulting deviation scale remains unreliable. Local calibration is therefore part of model application rather than a technical correction performed after the fact: the large healthy cohort defines the normative reference model, local observations anchor it within the target environment, and explicit transfer diagnostics determine whether individual regional scores remain interpretable (Bayer et al., 2022; Kia et al., 2022; Maccioni et al., 2026; Rutherford et al., 2022).

### 4.3 Amyloid-related molecular-enriched deviations in cognitively unimpaired participants

The second objective was to explore the transferred normative representation in a biologically relevant application: functional variation associated with amyloid accumulation before measurable cognitive impairment. The main result was not a diffuse increase in abnormality across monoaminergic systems. Instead, group differences were modest and selective, with the strongest corrected evidence involving DAT-enriched FC. The intermediate-amyloid-burden and amyloid-positive groups differed both in the subject-level burden of extreme DAT deviations and in the signed normative deviation of right caudal middle frontal cortex, whereas no regional NET- or SERT-enriched group difference survived correction. This selectivity is consistent with the intended role of region-wise NM: rather than assuming a homogeneous whole-brain response, the framework can identify which molecularly informed functional phenotypes depart from the expected age-related distribution. The primary subject-level outcome, RADB, was higher in the intermediate group in all four scopes, but none of these adjusted contrasts survived FDR-correction (DAT: *q* = 0.062). Normative-modeling studies of the Alzheimer continuum have likewise reported heterogeneous, individual-specific deviation patterns, including subtle deviations in non-demented individuals at risk (Kumar et al., 2025; Lawry Aguila et al., 2025; Verdi et al., 2023); the present results extend this approach to molecular-enriched functional phenotypes.

The right caudal middle frontal finding provides a spatially specific component to the subject-level DAT result. Because the DAT template is dominated by striatal signal and cortical dopamine clearance relies substantially on NET (Morón et al., 2002), a cortical DAT-enriched value reflects how strongly the regional BOLD signal is coupled to the predominantly striatal DAT-weighted fluctuation, not cortical DAT expression. The finding can therefore be read as a difference in the coupling of lateral prefrontal cortex with the dopaminergic system, consistent with the role of frontostriatal dopamine in cognitive control (Cools & D’Esposito, 2011). Previous molecular-imaging and molecular-atlas-informed studies have implicated dopaminergic alterations along the clinical Alzheimer continuum (Manca et al., 2025; Pilotto et al., 2025; Sala et al., 2021), thus reinforcing our results.

The direction of the categorical effects also argues against a simple interpretation in which molecular-enriched functional abnormality increases monotonically with global amyloid burden. Relative to the local amyloid-negative reference, the intermediate group showed lower DAT-enriched deviation scores in four right frontal and precentral regions, whereas in the right caudal middle frontal feature the amyloid-positive group lay at or above the reference location, and its DAT RADB was lower than that of the reference group. Intermediate-amyloid-burden and amyloid-positive participants are cross-sectional groups rather than successive observations of the same individuals. Functional network reorganization during early amyloid accumulation may nevertheless be stage-dependent, with phases of hyperconnectivity and hypoconnectivity and inverted U-shaped associations between amyloid and connectivity reported in cognitively normal older adults (Palmqvist et al., 2017; Schultz et al., 2017; Wu et al., 2026), and individuals with similar global amyloid burden can differ in tau pathology, reserve, and time since amyloid onset (Betthauser et al., 2022; Cabeza et al., 2018; Schultz et al., 2017). The present data therefore support a difference in the distribution of DAT-enriched functional deviations between amyloid groups, but do not establish whether that difference reflects vulnerability, compensation, adaptation, or a temporal stage of disease progression.

### 4.4 Continuous amyloid burden and APOE *ɛ*4

The categorical findings were not accompanied by corrected linear associations with continuous CL within cognitively unimpaired participants with CL ≥ 10. Likewise, APOE *ɛ*4 carrier status did not significantly modify the subject-level association between CL and deviation burden after correction. These negative findings constrain the interpretation of the categorical effects: the present data do not support a simple linear increase or decrease in molecular-enriched normative deviation across the intermediate-to-amyloid-positive range.

At the same time, the continuous analysis was specifically designed to test linear associations and therefore cannot exclude non-linear or stage-dependent relationships. The categorical results should not be used as evidence for such a trajectory in the absence of an explicit non-linear analysis. The absence of a corrected CL-by-APOE interaction should likewise be interpreted cautiously given the additional stratification and sample size required for interaction testing.

### 4.5 Limitations

Several limitations define the scope of the conclusions. First, the seven-dataset healthy reference cohort was intentionally heterogeneous in demographics and imaging protocols. Hierarchical modeling uses this heterogeneity to estimate broad normative variation, but the resulting reference distributions cannot be assumed to generalize equally across all ancestries, educational backgrounds, socioeconomic contexts, scanner environments, or clinical populations. In addition, amyloid status was not available for the older participants in the public normative datasets. Some cognitively healthy older individuals may therefore have harbored preclinical amyloid pathology, whose prevalence in cognitively normal adults rises steeply with age (Jansen et al., 2015), meaning that the reference trajectories should be interpreted as trajectories of clinically healthy aging rather than biomarker-confirmed amyloid-negative aging.

The local AMYPAD-PNHS reference sample introduces a related but distinct consideration. Although participants used for calibration were cognitively unimpaired and amyloid-negative, they were recruited through Alzheimer-focused AMYPAD parent cohorts rather than from a single unselected population-based sample (Pieperhoff et al., 2026). The included parent cohorts span research and clinically enriched recruitment settings; consequently, the local reference distribution may contain sources of variability that differ from those of a general-population sample. This does not prevent local calibration, but it should be considered when interpreting the resulting deviation scale and generalizing the biological findings. Because all amyloid-negative participants were used for local adaptation and calibration, no independent amyloid-negative comparison group was available, and the only fully independent categorical contrast was between the intermediate-amyloid-burden and amyloid-positive groups; future deployments could reserve a subset of eligible reference participants as an independent control group.

Second, REACT uses population-average molecular templates and therefore provides molecularly informed functional phenotypes rather than participant-specific measurements of neurotransmitter systems; it also assumes that the spatial distribution of each target is largely shared across participants. DAT, NET, and SERT were entered jointly in both REACT stages, so that shared BOLD variance was partitioned across them, although spatial overlap between templates, particularly the DAT and SERT, can still limit target specificity; this design is consistent with multitemplate approaches designed to reduce omitted-variable bias (Lawn et al., 2023); template collinearity was not re-estimated for the present analysis mask, but the three templates are a subset of the six-template set for which Lawn et al. (2024) reported variance-inflation factors below 5, and removing templates from a joint model cannot increase the variance-inflation factors of those retained. Individual molecular imaging would be required to establish whether a REACT-derived functional deviation co-varies with local transporter availability in the same participant.

Third, the normative feature space was restricted to the 68 cortical parcels of the DK atlas. Regional averaging improves robustness and preserves a common feature definition across datasets, but it can obscure more localized effects and excludes subcortical and cerebellar structures. Extending the framework to those regions is a logical next step but will require careful validation of registration and resting-state signal quality.

Fourth, the analytical cohort was restricted to cognitively unimpaired participants, operationally defined as having a global CDR score of 0. This restriction is useful for examining amyloid-related functional variation before overt dementia, but it limits the range of contemporaneous clinical impairment and reduces the ability to link deviations to cognitive outcomes. The present study therefore addresses variation in a clinically early group defined by global CDR = 0 rather than the full clinical Alzheimer continuum. Head motion was addressed during preprocessing but was not included as a covariate in the amyloid-group analyses; because in-scanner motion can bias functional connectivity estimates even after denoising, residual motion differences between groups cannot be fully excluded (Ciric et al., 2018; Power et al., 2012).

Fifth, global CL represents fibrillar amyloid burden rather than the complete Alzheimer biological state. Regional amyloid, tau, neurodegeneration, inflammation, cerebrovascular injury, and reserve were not modeled jointly. The operational CL = 10 and CL = 30 boundaries are useful analytical strata but should not be interpreted as direct transitions in monoaminergic physiology. This biological heterogeneity is one reason why a spatially resolved normative representation may be informative, but it also limits mechanistic interpretation of the present group differences.

Finally, the amyloid application was cross-sectional and exploratory. Intermediate-amyloid-burden and amyloid-positive groups are not successive observations of the same individuals, so the data cannot determine whether the observed deviations precede, increase with, compensate for, or follow other Alzheimer-related biological changes. Longitudinal imaging will be required to determine whether individual deviation profiles predict subsequent biomarker or cognitive change. Spatially resolved follow-up analyses could additionally integrate imaging transcriptomics to test whether regional molecular-enriched deviation patterns align with gene-expression profiles implicated in amyloid metabolism, neuroinflammation, or synaptic remodeling, as recently demonstrated for amyloid-related structure–function changes in AMYPAD-PNHS (Arunachalam et al., 2026; Martins et al., 2021).

## 5 Conclusion

This study establishes a region-wise normative framework for molecular-enriched functional connectivity and demonstrates that such models can be deployed beyond the datasets used for their original estimation when external application is accompanied by local hierarchical adaptation, calibration, and explicit feature-level diagnostics. This addresses the primary methodological objective of the work and provides a practical basis for distributing REACT-derived normative reference models across independently processed neuroimaging environments rather than requiring centralized reprocessing of all target data.

The exploratory AMYPAD-PNHS application further shows how this framework can be used to characterize individualized functional variation during cognitively unimpaired stages of amyloid accumulation. The observed effects were modest and selective rather than widespread, emphasizing the value of retaining regional and molecular-system information instead of reducing functional heterogeneity to a single global score.

## Acknowledgments

The authors thank all participants, investigators, and data-management teams who contributed to the datasets used in this study.

Data collection and sharing for the Cambridge Centre for Ageing and Neuroscience (Cam-CAN) were supported by the UK Biotechnology and Biological Sciences Research Council (grant BB/H008217/1), together with support from the UK Medical Research Council and the University of Cambridge, UK.

The Dallas Lifespan Brain Study (DLBS) was supported by the National Institutes of Health, National Institute on Aging grants 5R37AG-006265-27 and RC1AG036199 to Denise C. Park. The authors thank the DLBS participants for their contribution to this resource.

Research reported using the HCP-Aging data was supported by the National Institute on Aging of the National Institutes of Health under Award U01AG052564 and by the McDonnell Center for Systems Neuroscience at Washington University in St. Louis. The HCP-Aging 2.0 Release data used in this work came from DOI: 10.15154/1520707.

Data were provided in part by the Human Connectome Project, WU-Minn Consortium (Principal Investigators: David Van Essen and Kamil Ugurbil; 1U54MH091657), funded by the 16 NIH Institutes and Centers that support the NIH Blueprint for Neuroscience Research and by the McDonnell Center for Systems Neuroscience at Washington University.

The UCLA Consortium for Neuropsychiatric Phenomics dataset was supported by NIH Roadmap for Medical Research grants UL1-DE019580, RL1MH083268, RL1MH083269, RL1DA024853, RL1MH083270, RL1LM009833, PL1MH083271, and PL1NS062410. Preparation of the shared data in BIDS format was supported by the Laura and John Arnold Foundation.

Data collection and sharing for the Enhanced Nathan Kline Institute–Rockland Sample were supported by National Institute of Mental Health grant R01MH094639-01 and additional support from the New York State Office of Mental Health, the Research Foundation for Mental Hygiene, the Child Mind Institute (1FDN2012-1), the Center for the Developing Brain at the Child Mind Institute, National Institute of Mental Health grants R01MH081218, R01MH083246, and R21MH084126, the NKI Center for Advanced Brain Imaging, the Brain Research Foundation, and the Stavros Niarchos Foundation.

The Southwest University Adult Lifespan Dataset was supported by the National Natural Science Foundation of China (31571137 and 31500885), the National Outstanding Young People Plan, the Program for Top Young Talents by Chongqing, the Fundamental Research Funds for the Central Universities (SWU1509383, SWU1509451, and SWU1609177), the Natural Science Foundation of Chongqing (cstc2015jcyjA10106), and the Fok Ying Tung Education Foundation (151023).

Data used in the target-cohort analyses were obtained from the Prognostic and Natural History Study (PNHS), provided by the Amyloid Imaging to Prevent Alzheimer’s Disease (AMYPAD) Consortium. Investigators within AMYPAD-PNHS and the AMYPAD Consortium contributed to the design and implementation of AMYPAD and/or provided data but did not participate in the present analysis unless listed as authors. AMYPAD-PNHS received funding from the Innovative Medicines Initiative 2 Joint Undertaking under grant agreement No. 115952, with support from the European Union’s Horizon 2020 research and innovation programme and EFPIA (Pieperhoff et al., 2026).

## Data and Code Availability

The normative reference data are available from their original repositories under the corresponding access procedures and data-use conditions for Cam-CAN, DLBS, HCP-Aging, HCP-YA, UCLA CNP/LA5c, NKI-RS, and SALD. AMYPAD-PNHS data are available under controlled access through the Alzheimer’s Disease Data Initiative Workbench, as described by Pieperhoff et al. (2026). REACT is openly available from the react-fmri repository (https://github.com/ottaviadipasquale/react-fmri), and PCNtoolkit is openly available at https://github.com/predictive-clinical-neuroscience/PCNtoolkit and documented at https://pcntoolkit.readthedocs.io/ (S. de Boer et al., 2026).

Study-specific scripts for normative-model specification and transfer, feature-level QC, and the statistical analyses reported here will be made available from the corresponding author upon reasonable request and subject to the data-use restrictions of the contributing datasets.

## Author Contributions

Marco Pinamonti: Conceptualization, Methodology, Software, Formal analysis, Investigation, Visualization, Data curation, Writing – original draft. Manuela Moretto: Conceptualization, Methodology, Supervision, Writing – review and editing. Leonard Pieperhoff: Resources, Data curation, Methodology, Writing – review and editing. Prithvi Arunachalam: Methodology, Data curation, Software, Writing – review and editing. Frederik Barkhof: Resources, Data curation, Writing – review. Alle Meije Wink: Resources, Data curation, Software, Writing – review. Luigi Lorenzini: Conceptualization, Methodology, Supervision, Resources, Writing – review and editing. Mattia Veronese: Conceptualization, Methodology, Supervision, Project administration, Funding acquisition, Writing – review and editing.

## A Supplementary Materials

### A.1 Image processing

#### A.1.1 MRI preprocessing and quality control

##### Healthy normative reference cohort

The raw T1w and rs-fMRI data from all seven normative reference datasets were processed using an identical workflow to minimize methodological variability across cohorts. The organization of imaging data followed the Brain Imaging Data Structure (BIDS) specification (Gorgolewski et al., 2016), and the data were processed using standardized, BIDS-compatible neuroimaging tools. The degree to which the BIDS specification was adhered to was determined by means of the BIDS Validator (v1.14.6) prior to image processing (Blair et al., 2024).

##### Quality control

The assessment of image quality was conducted by employing the MRIQC (v24.1.0) tool (Esteban et al., 2017), with each dataset and imaging modality undergoing independent evaluation. Given the substantial number of scans included in the healthy normative reference cohort, QC followed a semi-automated procedure combining quantitative image-quality metrics (IQMs) with targeted visual inspection. Group-level MRIQC reports were examined separately within each dataset, and scans with IQMs falling outside the inner Tukey fences, defined as values below *Q*_1_ − 1.5 × IQR or above *Q*_3_ + 1.5 × IQR, were flagged for subsequent visual assessment (Tukey, 1977).

For T1w images, the quality assessment focused on signal-to-noise ratio (SNR), gray-to-white-matter contrast-to-noise ratio, and the entropy-focus criterion. For rs-fMRI data, temporal signal-to-noise ratio (tSNR), mean framewise displacement (FD), and the temporal derivative of the variance of the BOLD signal (DVARS) were considered. All scans that were flagged by the quantitative procedure were then subjected to a visual inspection. Images that exhibited substantial motion, ringing, cropping, or other evident acquisition artifacts were excluded from further processing.

##### Structural and functional preprocessing with fMRIPrep

Preprocessing of the structural and fMRI data was conducted using fMRIPrep (v23.2.2) (Esteban et al., 2019, 2020). For each participant, T1w images underwent intensity non-uniformity correction using N4 bias-field correction, implemented in Advanced Normalization Tools (ANTs) (v2.4.4) (Tustison et al., 2010). The skull stripping procedure was executed employing the Nipype implementation of antsBrainExtraction.sh which utilized the OASIS30ANTs template distributed through TemplateFlow (v23.1.0) (Ciric et al., 2022). The brain-extracted T1w image was segmented into CSF, WM, and gray matter (GM) using FAST from FSL (v6.0.6.2) (Zhang et al., 2001). Cortical surface reconstruction was performed using recon-all from FreeSurfer (v7.3.2) (Fischl, 2012), and the initial brain mask was refined by reconciling the ANTs-and FreeSurfer-derived gray-matter segmentations using a Mindboggle-based procedure (Klein et al., 2017). Subsequently, the anatomical images were nonlinearly normalized to the MNI ICBM 152 nonlinear sixth-generation asymmetric template (MNI152NLin6Asym) at 2-mm isotropic resolution (Ciric et al., 2022; Evans et al., 2012). Nonlinear registration was performed using antsRegistration from ANTs (Avants et al., 2008, 2011).

For each rs-fMRI acquisition, a BOLD reference image was generated and head-motion parameters were estimated using mcflirt from FSL before temporal filtering (Jenkinson et al., 2012). Subsequent to this, slice-timing correction was performed, and susceptibility-distortion correction was applied when the required fieldmap information was available. The BOLD reference image was co-registered with the corresponding T1w image using FreeSurfer’s boundary-based registration (bbregister) with six degrees of freedom (Greve & Fischl, 2009). A series of transformations were concatenated and applied in a single interpolation step using nitransforms with cubic B-spline interpolation. These transformations included head motion, BOLD-to-T1w, and T1w-to-MNI spatial transformations.

Framewise displacement (FD) and DVARS were retained as framewise motion-sensitive quality metrics (Power et al., 2012). For the present preprocessing workflow, volumes exceeding either FD > 0.5 mm or standardized DVARS > 1.5 were classified as motion-affected frames for subsequent interpolation. No nuisance regression or temporal filtering was applied within fMRIPrep.

##### Post-processing with XCP-D

The fMRIPrep derivatives were subsequently denoised using XCP-D (v0.7.4) (Mehta et al., 2024). Confound regression was executed in accordance with the acompcor nuisance-regression strategy, encompassing the six rigid-body motion parameters and their temporal derivatives, in conjunction with the ten leading anatomical CompCor (aCompCor) components derived from WM and CSF (Behzadi et al., 2007; Satterthwaite et al., 2013).

Frames that were identified as being affected by motion were reconstructed using cubic-spline interpolation to ensure the preservation of a temporally continuous BOLD signal for the purpose of subsequent REACT analysis. The BOLD time series and nuisance regressors underwent temporal high-pass filtering using a second-order Butterworth filter with a cutoff frequency of 0.01 Hz. Subsequently, the nuisance effects were eliminated through the implementation of linear regression with the specified confound set and cosine basis functions. This process followed established methodologies for the mitigation of motion-related artifacts in FC analyses (Ciric et al., 2018). Given that REACT necessitates continuous functional time series, the interpolated, non-censored residual BOLD images generated by XCP-D were preserved for subsequent processing. No spatial smoothing was applied within XCP-D.

##### Spatial smoothing

Following XCP-D post-processing, the interpolated residual BOLD images were spatially smoothed using FSL (v6.0.6.2) with an isotropic Gaussian kernel of 6 mm FWHM, as described in Jenkinson et al. (2012) and Smith et al. (2004). The resulting denoised and spatially smoothed BOLD images constituted the functional input to the subsequent REACT analysis.

##### AMYPAD-PNHS cohort

The rs-fMRI data from the AMYPAD-PNHS cohort had previously undergone a standardized preprocessing workflow, as described by Arunachalam et al. (2026). The preprocessing of functional images was conducted using fMRIPrep (v23.0.1). In summary, the preprocessing stage entailed the estimation of a BOLD reference image, the extraction of head-motion parameters, and the implementation of slice-timing correction when applicable. Corrections for susceptibility-related distortions were implemented through the utilization of a fieldmap-less approach, which was based on the non-linear registration of the BOLD reference image to the corresponding T1w image. This registration was accomplished through the implementation of symmetric normalization in ANTs. Subsequent to this, the functional images were aligned to the T1w image using FreeSurfer’s boundary-based registration and transformed with ANTs to MNI152NLin6Asym standard space at 2-mm isotropic resolution, matching the space used for the molecular templates and regional feature-generation workflow.

Thereafter, the initial non-steady-state volumes were removed following fMRIPrep preprocessing. The functional time series were spatially smoothed using a Gaussian kernel with a FWHM of 4 millimeters. The nuisance regression model incorporated six rigid-body head-motion parameters, along with the mean cerebrospinal-fluid and white-matter signals. Linear trends were removed, and the resulting time series were temporally band-pass filtered between 0.01 and 0.1 Hz. The resulting preprocessed BOLD images constituted the functional input to the molecular-enriched connectivity analyses performed in the present study.

From this stage onward, the AMYPAD-PNHS and healthy normative-reference cohorts underwent the same molecular-enriched feature-generation workflow. Specifically, REACT was applied to the AMYPAD-PNHS data using the same DAT, NET, and SERT molecular templates, analysis masks, and two-stage regression procedure described in Section A.1.2. The resulting subject-specific molecular-enriched FC maps were then parcellated using the same anatomical definitions and summarized using the same regional mean-extraction procedure described in Section A.1.2. This common downstream feature-definition strategy yielded nominally matched regional molecular-enriched functional-connectivity variables in the normative-reference and AMYPAD-PNHS cohorts, providing the feature correspondence required for normative-model transfer while leaving upstream acquisition and preprocessing differences to be addressed by hierarchical adaptation and local calibration.

The use of independently established upstream preprocessing workflows was deliberate and formed part of the transferability evaluation. AMYPAD-PNHS raw data were not retrospectively reprocessed with the complete normative-reference preprocessing pipeline, and no explicit feature-level harmonization was applied before normative-model transfer. Instead, cross-cohort compatibility was imposed at the molecular-enriched feature-definition stage by applying the same REACT templates, analysis masks, cortical parcellation, and regional extraction procedure. Hierarchical transfer and local calibration were then used to test whether the pretrained NM could accommodate the target-domain distribution under this deployment-oriented scenario.

#### A.1.2 Molecular-enriched functional connectivity

##### Receptor-Enriched Analysis of functional Connectivity by Targets

Molecular-enriched FC was derived using REACT (Dipasquale et al., 2019). REACT integrates individual rs-fMRI BOLD fluctuations with population-level molecular density templates obtained using PET or SPECT.

The analysis comprised two consecutive multivariate general linear models (GLMs). In Stage 1, the DAT, NET, and SERT molecular-density templates were entered jointly in a single spatial design matrix and regressed against each participant’s BOLD data. This yielded three subject-specific target-weighted time series while accounting for spatial covariance among the molecular templates. Stage 1 was restricted to the intersection of the common positive support of the three molecular templates, which were supplied together as a single four-dimensional atlas, and the gray-matter mask distributed with the REACT implementation. In Stage 2, the three Stage-1 time series were entered jointly into a temporal regression against the voxel-wise BOLD signal within the gray-matter analysis mask, yielding one subject-specific molecular-enriched FC map for each molecular target. Joint inclusion of molecular maps is important when templates share spatial variance, because omission of correlated molecular systems can alter target-specific attribution of BOLD variance (Lawn et al., 2023); previous multitemplate REACT work that included the present DAT, NET, and SERT templates evaluated this issue using spatial correlations and variance-inflation diagnostics (Lawn et al., 2024). The same joint design, masks, and template definitions were used for the normative-reference and AMYPAD-PNHS cohorts.

##### Molecular templates

Three monoaminergic transporter templates were included: the DAT, NET, and SERT. The DAT template was derived from [^123^I]-ioflupane SPECT data from 26 subjects with no evidence of nigrostriatal degeneration (García-Gómez et al., 2013). The NET template was generated from [^11^C]MRB PET parametric maps acquired from 10 healthy individuals (Hesse et al., 2017). The SERT template was derived from the [^11^C]DASB PET component of a high-resolution serotonin atlas built from 210 healthy individuals aged 18–45 years (Beliveau et al., 2017). All templates were represented in MNI152NLin6Asym space at 2-mm isotropic resolution and scaled between 0 and 1.

##### Regional feature extraction

Subject-specific DAT-, NET-, and SERT-enriched FC maps were summarized using the DK cortical parcellation. For each molecular system, the mean molecular-enriched FC value was extracted from 34 DK regions in each hemisphere. These regions included the banks of the superior temporal sulcus, caudal anterior cingulate, caudal middle frontal, cuneus, entorhinal, fusiform, inferior parietal, inferior temporal, isthmus cingulate, lateral occipital, lateral orbitofrontal, lingual, medial orbitofrontal, middle temporal, parahippocampal, paracentral, pars opercularis, pars orbitalis, pars triangularis, pericalcarine, postcentral, posterior cingulate, precentral, precuneus, rostral anterior cingulate, rostral middle frontal, superior frontal, superior parietal, superior temporal, supramarginal, frontal pole, temporal pole, transverse temporal, and insula. This yielded 68 cortical measurements for each molecular system and 204 cortical molecular-system features in total (68 DAT, 68 NET, and 68 SERT). A separate normative model was estimated for each cortical molecular-system feature. For the AMYPAD-PNHS cohort, subject-specific FreeSurfer DK labels were those produced by the standard AMYPAD structural processing workflow, which uses FreeSurfer v7.1.1 for cortical reconstruction and parcellation. For regional extraction, subject-specific FreeSurfer DK labels were propagated from FreeSurfer native anatomical space to T1w space and then to MNI152NLin6Asym space by combining the subject-specific FreeSurfer-native-to-T1w registration with the fMRIPrep T1w-to-MNI152NLin6Asym transform. Label images were resampled with nearest-neighbor interpolation to preserve discrete parcel identities before regional averaging of the REACT maps.

### A.2 Sensitivity analyses

Sensitivity analyses evaluated whether the principal findings depended on model-diagnostic filtering, deviation-score representation, downstream covariate adjustment, or RADB distributional handling. First, the principal analyses were repeated in the strict cross-batch diagnostic sensitivity domain containing only cortical features that satisfied both the normative-model and transfer-diagnostic criteria in all three AMYPAD acquisition batches. Regional multiplicity correction in this sensitivity analysis was applied across the retained diagnostic feature domain, while subject-level FDR correction remained across the four global/system-specific scopes.

Second, independent intermediate-versus-positive analyses were repeated using model-native *Z*^raw^. Because this score can retain residual batch-specific location and scale differences, regional and RADB effects were estimated separately within each acquisition batch using HC3 regression adjusted for age and sex. Each batch-specific coefficient and SE was divided by the corresponding model residual SD to obtain a standardized effect, and standardized effects were combined across batches using inverse-variance fixed-effect meta-analysis. Cochran’s *Q*and *I*^2^ characterized between-batch heterogeneity. The same batch-stratified strategy was used for *Z*^raw^ continuous CL analyses. FDR correction followed the same 204-feature regional or four-scope subject-level families used in the corresponding atlas-complete primary analyses.

For *Z*^raw^ extreme-deviation burden, a universal |*Z*| > 1.96 threshold was not applied because *Z*^raw^ could retain batch-dependent scaling. Instead, feature- and batch-specific empirical 2.5th and 97.5th percentile limits were estimated from the corresponding local amyloid-negative reference participants, and *Z*^raw^ extreme-deviation burden was defined as the proportion of features falling outside these empirical local 95% limits.

RADB distributional sensitivity analyses flagged values beyond a 3×IQR outer fence within each global or molecular-system-specific scope; primary analyses retained these observations, while sensitivity models were repeated after their exclusion. RADB group analyses were also repeated after natural-log transformation.

#### A.2.1#Effects of alternative analytical specifications

Restricting the analyses to the 157 cortical features that satisfied both normative and transfer diagnostic criteria in all three AMYPAD batches yielded adjusted intermediate minus amyloid-positive RADB differences of 0.169 for the global scope, 0.153 for DAT, 0.086 for NET, and 0.164 for SERT; none survived FDR correction (global, DAT, and SERT *q* = 0.063; NET *q* = 0.283). The DAT extreme-deviation-burden difference was 0.029 (95% CI 0.004–0.054, *p* = 0.022, *q* = 0.090). Within this strict diagnostic sensitivity domain, no tested regional intermediate versus amyloid-positive effect or continuous CL association survived FDR correction.

In the independent model-native *Z*^raw^ intermediate versus amyloid-positive analysis, DAT-enriched right caudal middle frontal cortex again showed the strongest regional group effect. The pooled standardized effect was −0.698 (95% CI [−1.036, −0.361], *p* < 0.001, *q* = 0.010), with no evidence of between-batch heterogeneity (*I*^2^ = 0%). Batch-specific standardized effects were −0.689 in BBRC-ALFA, −0.630 in FACEHBI, and −0.758 in VUmc.

For subject-level RADB calculated from *Z*^raw^, batch-stratified fixed-effect meta-analysis yielded a global intermediate minus amyloid-positive standardized effect of 0.369 (95% CI 0.082–0.656, *p* = 0.012, *q* = 0.047), with *I*^2^ = 53.6%. System-specific *Z*^raw^ RADB effects did not survive the four-scope correction (DAT *q* = 0.172, NET *q* = 0.060, SERT *q*= 0.054). Using feature- and batch-specific empirical local-reference 95% limits for *Z*^raw^ extreme deviations, intermediate-burden minus amyloid-positive differences were 0.026 for global (*q* = 0.044), 0.034 for DAT (*q* = 0.033), 0.009 for NET (*q* = 0.589), and 0.036 for SERT (*q* = 0.044). Neither *Z*^raw^ regional nor subject-level continuous CL analyses yielded statistically significant associations after FDR corrections.

Fourteen scope-specific RADB values from nine participants were flagged by the 3×IQR outer-fence rule and were retained in the primary analyses. Excluding these values preserved positive intermediate minus amyloid-positive RADB coefficients across all four scopes, with no comparison surviving FDR correction (minimum *q*= 0.141). Natural-log transformation of RADB likewise yielded no significant group comparison after FDR correction (minimum *q*= 0.117).

The robustness of the regional intermediate amyloid burden versus amyloid-positive group coefficients to downstream covariate adjustment is illustrated in Supplementary Figure 8.

#### A.2.2#Robustness of the primary findings

The sensitivity analyses showed that the DAT-enriched right caudal middle frontal group difference was consistent in direction and magnitude across covariate specifications, score representations, and acquisition batches (batch-specific standardized effects −0.63 to −0.76; *I*^2^ = 0%). However, the corrected categorical findings were attenuated when the analysis was restricted to the most conservative 157-feature cross-batch diagnostic domain. Thus, the main regional finding was robust to downstream adjustment and deviation-score representation but remained sensitive to the definition of feature-level transfer quality. Independent replication with prospectively defined transfer criteria will be required to determine its generalizability.

**Figure 8:**
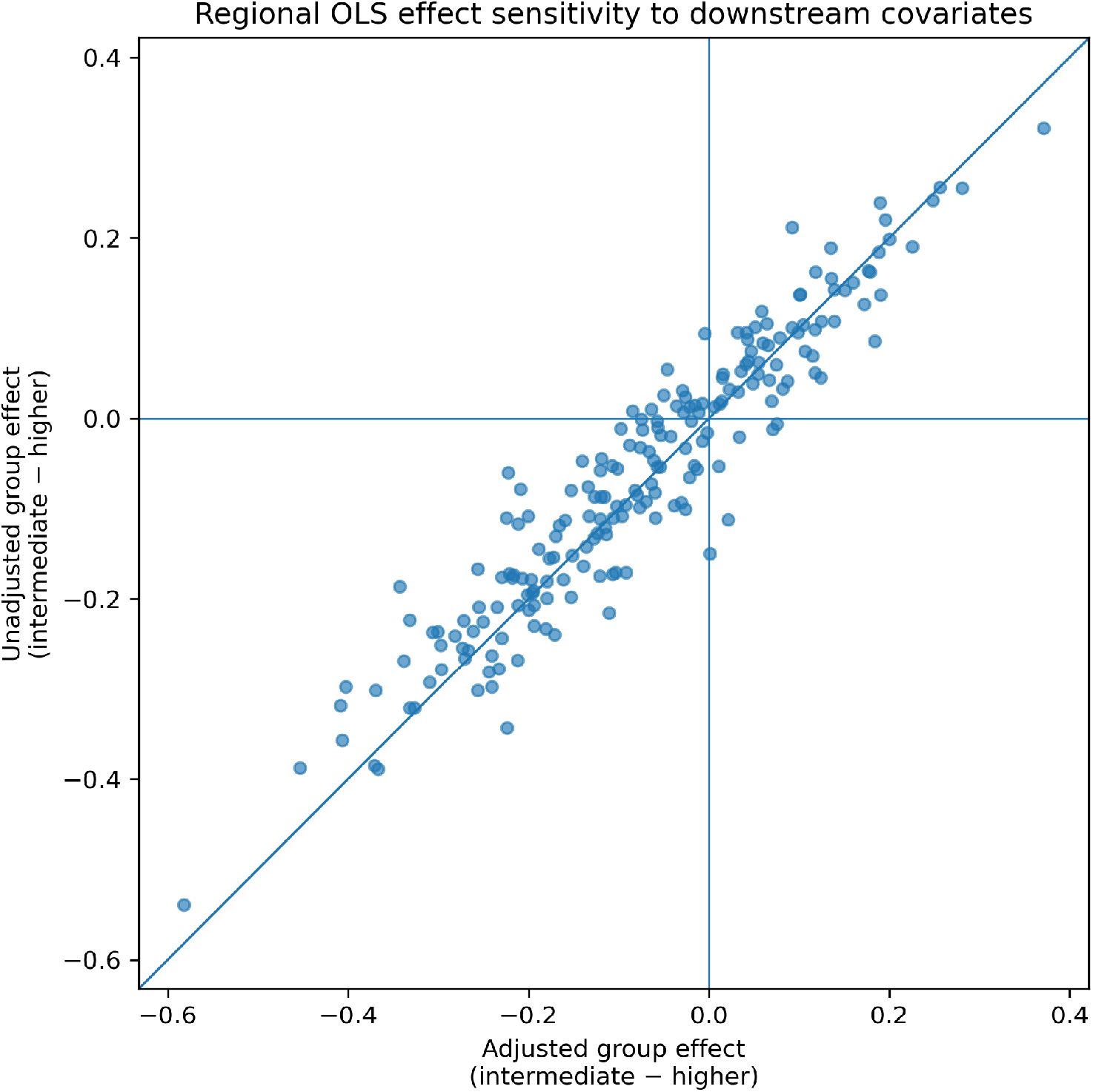
Sensitivity of regional intermediate minus amyloid-positive group coefÏcients to downstream covariate adjustment. Each point represents one cortical molecular-system feature. The horizontal axis shows the HC3 coefÏcient from the primary model adjusted for age, sex, and acquisition batch; the vertical axis shows the coefÏcient from the group-only HC3 model. The diagonal indicates equality of the two estimates.

